# Clustering of adult-onset diabetes into novel subgroups guides therapy and improves prediction of outcome

**DOI:** 10.1101/186387

**Authors:** E Ahlqvist, P Storm, A Käräjämäki, M Martinell, M Dorkhan, A Carlsson, P Vikman, RB Prasad, D Mansour Aly, P Almgren, Y Wessman, N Shaat, P Spegel, H Mulder, E Lindholm, O Melander, O Hansson, U Malmqvist, Å Lernmark, K Lahti, T Forsén, T Tuomi, AH Rosengren, L Groop

## Abstract

**Background:** Diabetes is presently classified into two main forms, type 1 (T1D) and type 2 diabetes (T2D), but especially T2D is highly heterogeneous. A refined classification could provide a powerful tool individualize treatment regimes and identify individuals with increased risk of complications already at diagnosis.

**Methods:** We applied data-driven cluster analysis (k-means and hierarchical clustering) in newly diagnosed diabetic patients (N=8,980) from the Swedish ANDIS (All New Diabetics in Scania) cohort, using five variables (GAD-antibodies, BMI, HbA1c, HOMA2-B and HOMA2-IR), and related to prospective data on development of complications and prescription of medication from patient records. Replication was performed in three independent cohorts: the Scania Diabetes Registry (SDR, N=1466), ANDIU (All New Diabetics in Uppsala, N=844) and DIREVA (Diabetes Registry Vaasa, N=3485). Cox regression and logistic regression was used to compare time to medication, time to reaching the treatment goal and risk of diabetic complications and genetic associations.

**Findings:** We identified 5 replicable clusters of diabetes patients, with significantly different patient characteristics and risk of diabetic complications. Particularly, individuals in the most insulin-resistant cluster 3 had significantly higher risk of diabetic kidney disease, but had been prescribed similar diabetes treatment compared to the less susceptible individuals in clusters 4 and 5. The insulin deficient cluster 2 had the highest risk of retinopathy. In support of the clustering, genetic associations to the clusters differed from those seen in traditional T2D.

**Interpretation:** We could stratify patients into five subgroups predicting disease progression and development of diabetic complications more precisely than the current classification. This new substratificationn may help to tailor and target early treatment to patients who would benefit most, thereby representing a first step towards precision medicine in diabetes.

**Funding:** The funders of the study had no role in study design, data collection, analysis, interpretation or writing of the report.

**Research in context:** **Evidence before this study**

The current diabetes classification into T1D and T2D relies primarily on presence (T1D) or absence (T2D) of autoantibodies against pancreatic islet beta cell autoantigens and age at diagnosis (earlier for T1D). With this approach 75-85% of patients are classified as T2D. A third subgroup, Latent Autoimmune Diabetes in Adults (LADA,<10%), is defined by presence of autoantibodies against glutamate decarboxylase (GADA) with onset in adult age. In addition, several rare monogenic forms of diabetes have been described, including Maturity Onset Diabetes of the Young (MODY) and neonatal diabetes. This information is provided by national guidelines (ADA,WHO, IDF, Diabetes UK etc) but has not been much updated during the past 20 years and very few attempts have been made to explore heterogeneity of T2D. A topological analysis of potential T2D subgroups using electronic health records was published in 2015 but this information has not been implemented in the clinic.

**Added value of this study**

Here we applied a data-driven cluster analysis of 5 simple variables measured at diagnosis in 4 independent cohorts of newly-diagnosed diabetic patients (N=14755) and identified 5 replicable clusters of diabetes patients, with significantly different patient characteristics and risk of diabetic complications. Particularly, individuals in the most insulin-resistant cluster 3 had significantly higher risk of diabetic kidney disease.

**Implications of the available evidence**

This new sub-stratification may help to tailor and target early treatment to patients who would benefit most, thereby representing a first step towards precision medicine in diabetes

## Introduction

Diabetes is the fastest increasing disease worldwide and one of the greatest threats to human health.^1^ Unfortunately, current treatment strategies have not been able to stop the progressive course of the disease and prevent development of chronic diabetic complications. There are several explanations for these shortcomings. Diagnosis of diabetes is based upon measurement of only one metabolite, glucose, but the disease is very heterogeneous with regard to clinical presentation and progression.

The current diabetes classification into T1D and T2D relies primarily on presence (T1D) or absence (T2D) of autoantibodies against pancreatic islet beta cell autoantigens and age at diagnosis (earlier for T1D). With this approach 75-85% of patients are classified as T2D. A third subgroup, Latent Autoimmune Diabetes in Adults (LADA,<10%), defined by presence of autoantibodies against glutamate decarboxylase (GADA) is phenotypically indistinguishable from T2D at diagnosis but become more T1D-like with time.^2^ With the introduction of gene sequencing for clinical diagnostics several rare monogenic forms of diabetes were described, including Maturity Onset Diabetes of the Young (MODY) and neonatal diabetes.^3,4^

A limitation to current treatment guidelines is that they respond to poor metabolic control when it has developed but lack means to predict which patients will need intensified treatment. Importantly, accumulating evidence suggests that the early treatment is critical for prevention of life-shortening complications since target tissues seem to remember poor metabolic control decades later, also referred to as “metabolic memory”.^5,6^

A refined classification could provide a powerful tool to identify those at greatest risk of complications already at diagnosis, and enable individualized treatment regimes in the same way as a genetic diagnosis of monogenic forms of diabetes guides clinicians to optimal treatment.^7^ With this aim, we present a novel diabetes classification based on unsupervised data-driven cluster analysis of six commonly measured variables and compare it metabolically, genetically and clinically to the current classification in four separate populations from Sweden and Finland.

## Methods

### Study populations

**The ANDIS (All New Diabetics in Scania)** project (http://andis.ludc.med.lu.se/) aims to recruit all incident cases of diabetes within Scania (Skåne) County in southern Sweden, which has about 1,200,000 inhabitants. All health care providers in the region were invited; the current registration covered the period January 1^st^ 2008 until November 2016 during which 177 clinics registered 14,625 patients (> 90% of eligible patients), aged 0-96 years within a median of 40 days (IQR 12-99) after diagnosis.

**The Scania Diabetes Registry (SDR)** was recruited in the same region of Sweden between 1996 and 2009 including >7,400 individuals with diabetes of all types, 1,466 of whom were recruited no longer than two years after diagnosis and had all data necessary for clustering.^8^

**ANDIU (All new diabetics in Uppsala)** is a project similar to ANDIS started in 2011. Subjects are recruited in the Uppsala region (~300,000 inhabitants) in Sweden (http://www.andiu.se). N=844 patients had complete data for all clustering variables.

**DIREVA (Diabetes Registry Vaasa)** from Western Finland (~170,000 inhabitants) includes 5107 individuals with diabetes recruited 2009-2014.

**MDC-CVA (Malmö Diet and Cancer)** includes subjects living in Malmö, Southern Sweden who were invited to a clinical examination, including oral glucose tolerance tests to diagnose diabetes (n=3,300).^9^ Individuals without diabetes were used as controls for the genetic analyses.

### Measurements

In ANDIS blood samples were drawn at the registration visit for measurements of glucose, C-peptide, GAD autoantibodies, metabolites and DNA. Fasting plasma glucose was analyzed after an overnight fast using the HemoCue Glucose System (HemoCue AB, Ängelholm, Sweden). C-peptide concentrations were determined using an ElectroChemiLuminiscenceImmunoassay on Cobas e411 (Roche Diagnostics, Mannheim, Germany) or by. radioimmunoassay (Human C-peptide RIA; Linco, St Charles, MO, USA; or Peninsula Laboratories, Belmont, CA, USA). In ANDIS and SDR GADA was measured by an Enzyme-Linked Immunosorbent Assay (ELISA) (ref <11 U/ml^10^) or with radiobinding assays using 35S-labelled protein^11^ (positive cut-off: 5 RU or 32 IU/ml). The radiobinding assay showed 62–88% sensitivity and 91–99% specificity in workshops (Combinatorial Autoantibody or Diabetes/Islet Autoantibody Standardization Programs) from 1998 to 2013, and the ELISA assay showed 72% sensitivity and 99% specificity in the 2013 Islet Autoantibody Standardization Program workshop. In ANDIU GADA was measured at Laboratory Medicine in Uppsala (ref <5 U/ml). In DIREVA, GADA were measured with an ELISA assay (RSR, Cardiff, UK; positive cut-off 10 IU/ml). ZnT8A antibodies were measured using a Radio-Binding Assay as previously described.^12^ HbA1c was measured at diagnosis by the treating physician. HOMA2B and HOMA2IR were calculated from fasting C-peptide and glucose using the HOMA calculator (the University of Oxford, UK).^13^ Measurements of HbA1c, ALT, ketones and serum creatinine over time were obtained from the Clinical Chemistry department database.

### Genotyping

Genotyping was carried out using iPlex (Sequenom, San Diego, California, US) or TaqMan assays (Thermo Fisher Scientific). Patients and SNPs with call rate < 90% were excluded.

### Definitions of diabetic complications

The MDRD (Modification of Diet in Renal Disease) formula^14^ was used to calculate estimated glomerular filtration rate (eGFR). Chronic kidney disease (CKD) was defined as eGFR<60 (CKD stage 3A) or <45 (CKD stage 3B) for more than 90 days (onset of CKD was set as the start of the >90 day period). End-stage renal disease (ESRD) was defined as at least one eGFR below 15 mL/min/1·73m2.

Macroalbuminuria was defined as at least two out of three consecutive visits with albumin excretion rate (AER) ≥200 μg/min or AER ≥300 mg/24 h or albumin-creatinine ratio (ACR) ≥25/35 mg/mmol for men/women.

Diabetic retinopathy was diagnosed by an ophthalmologist based on fundus photographs.^15^ Coronary events (CE) were defined by ICD-10 codes I21, I252, I20, I251, I253, I254, I255, I256, I257, I258 and I259. Stroke was defined by ICD-10 codes I60, I61, I63 and I64.

### Cluster analysis

Cluster analysis was performed on scaled and centered values. Patients with secondary diabetes were excluded, as were extreme outliers (>5 SD). GADA was included as a dichotomous variable. TwoStep clustering was performed in SPSS v.23 for 2 to 15 clusters using log-likelihood as distance measure and Schwarz’s Bayesian criterion for clustering. K-means clustering was performed with k=4 using the kmeansruns function (runs=100) in the fpc package in R. Cluster centres in ANDIS used to classify patients in replication cohorts are presented in table s3. Assessment of clusterwise stability was carried out by resampling the dataset 2,000 times and computing the Jaccard similarities to the original cluster. Generally, a stable cluster should yield a Jaccard similarity >0·75.^16^

### Statistical analysis

Risk of complications was calculated using cox regression in SPSS v.23. Sex was included as a covariate in all analyses. Analysis including sex and age at onset of diabetes or first eGFR was also performed as indicated in the text.

Associations between clusters and genotypes were calculated using the MLE method in SNPtest2. The equality of odds ratios across strata in table1 was tested using seemingly unrelated estimation (suest) as implemented in Stata. Non-diabetic individuals from the MDC-CVA cohort were used as controls. Patients of known non-Swedish origin were excluded.

**Table 1.**
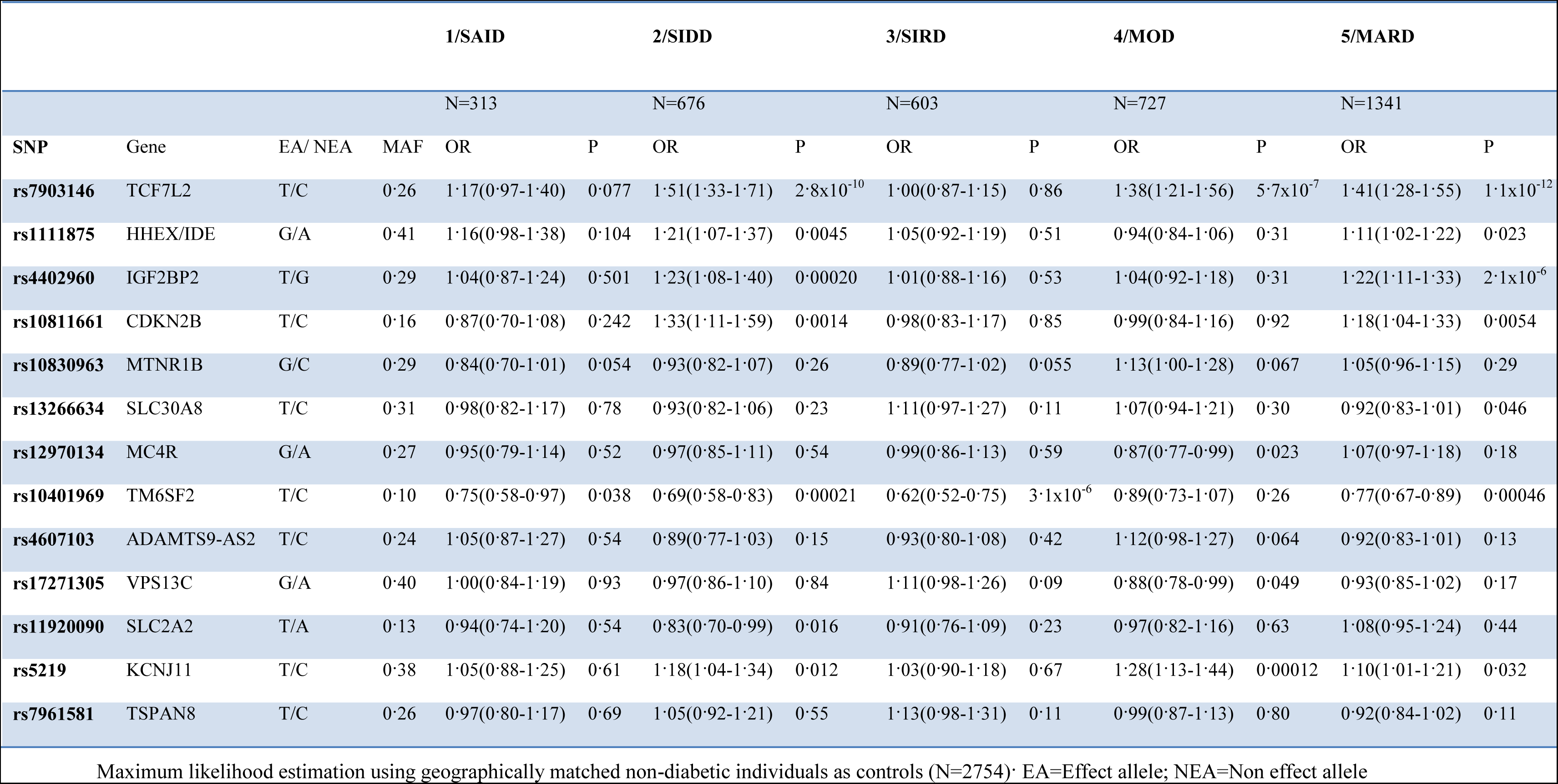
Genetic associations with specific ANDIS clusters reaching at least nominal significance for difference between clusters 2 to 5.

### Funding

The funding agencies had no role in study design, data collection, data analysis, data interpretation, or writing of the report.

## Results

We first analyzed a cohort of 14,652 newly diagnosed diabetic patients from Sweden termed ANDIS (All New Diabetics in Scania). Of them, 932 (6·4%) were registered before age 18 and not included in analyses of adult diabetes. Of the adult patients, 1·4% had T1D (defined as GADA positive and C-peptide < 0·3 nmol/l), 4·9% LADA (GADA-positive and C-peptide ≥ 0·3 nmol/l), 1·1% secondary diabetes (coexisting pancreatic disease) and 3·5% were unclassifiable due to missing data. The remaining patients (82·7%) were considered to have T2D (Table S1). Five quantitative variables (age, BMI, HbA1c, Homeostasis Model Assessment 2 estimates of beta-cell function [HOMA2-B] and insulin resistance [HOMA2-IR]), plus presence or absence of GADA as a binary variable, were used in a cluster analysis to reclassify patients into novel diabetes subgroups. Fasting C-peptide and plasma glucose were used to estimate HOMA2.^13^ Patients with complete data for the clustering variables (N=8,980) were included in the further analyses (Figure 1A, Table S1).

**Figure 1.**
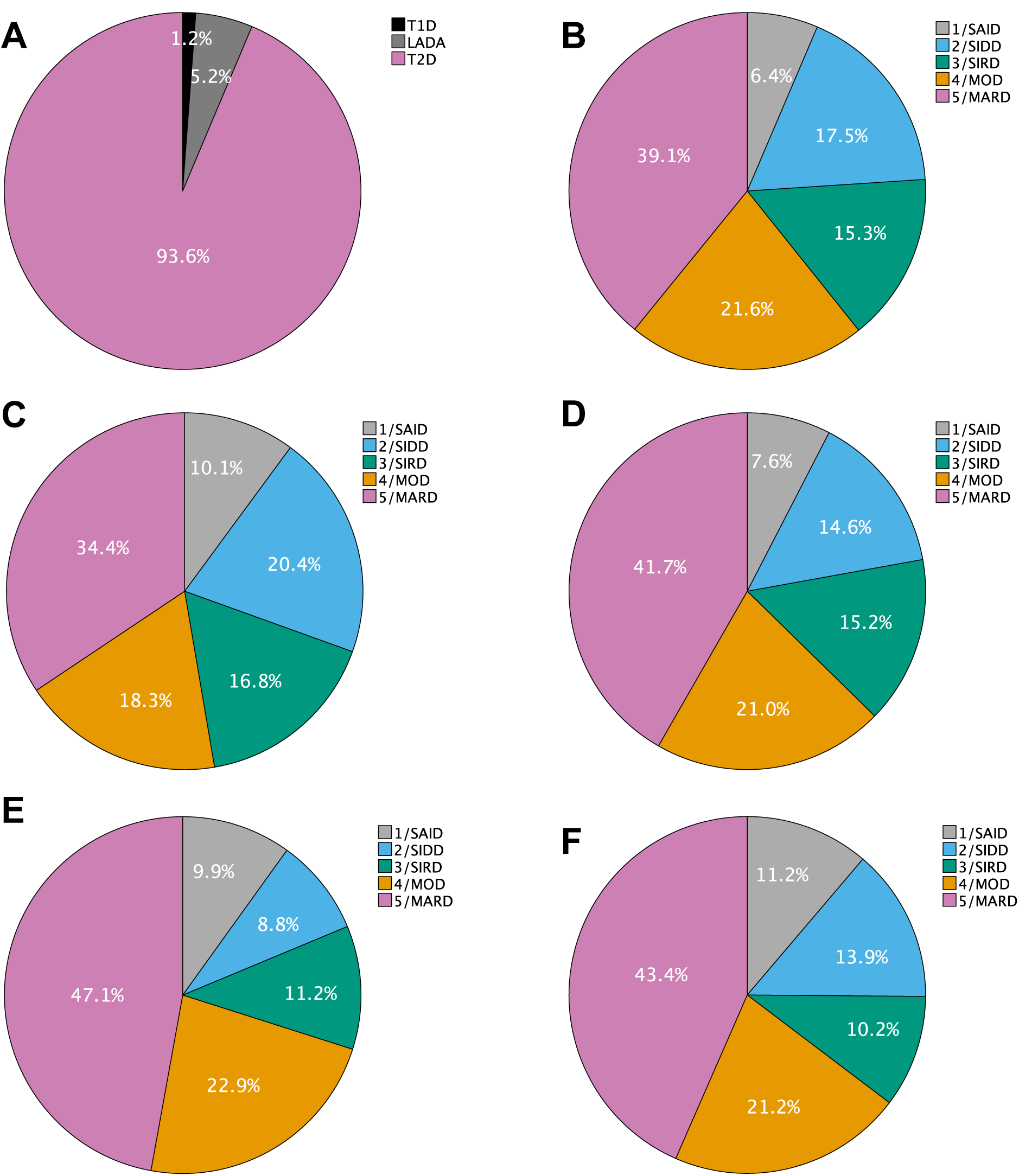
Patient distribution using different methods for classification. Distribution of ANDIS patients included in the clustering using (A) traditional classification and (B) k-means clustering. Distribution of patients using k-means clustering in SDR (C), ANDIU (D) and in DIREVA stratified for newly diagnosed (E) and long duration (F).

First, we applied TwoStep clustering; the first step estimates the optimal number of clusters based upon silhouette width and the second step performs hierarchical clustering. Men (N=5,334) and women (N=3,646) were clustered separately; patients with secondary diabetes were excluded from subsequent analyses. The minimum silhouette width was found for 5 clusters in both men and women, exhibiting similar cluster distributions and characteristics (Figure S1). We verified the results using a second clustering method, k-means clustering. Because all GADA-positive patients clustered together in TwoStep analysis, and k-means clustering is inappropriate for binary variables, we restricted the k-means clustering to GADA-negative patients. This resulted in a similar cluster distribution as TwoStep with the same overall cluster characteristics in both sexes (Figure 1B, 2 and S2). Cluster stability was estimated as Jaccard means, which were >0·8 for all clusters regardless of sex.

**Figure 2.**
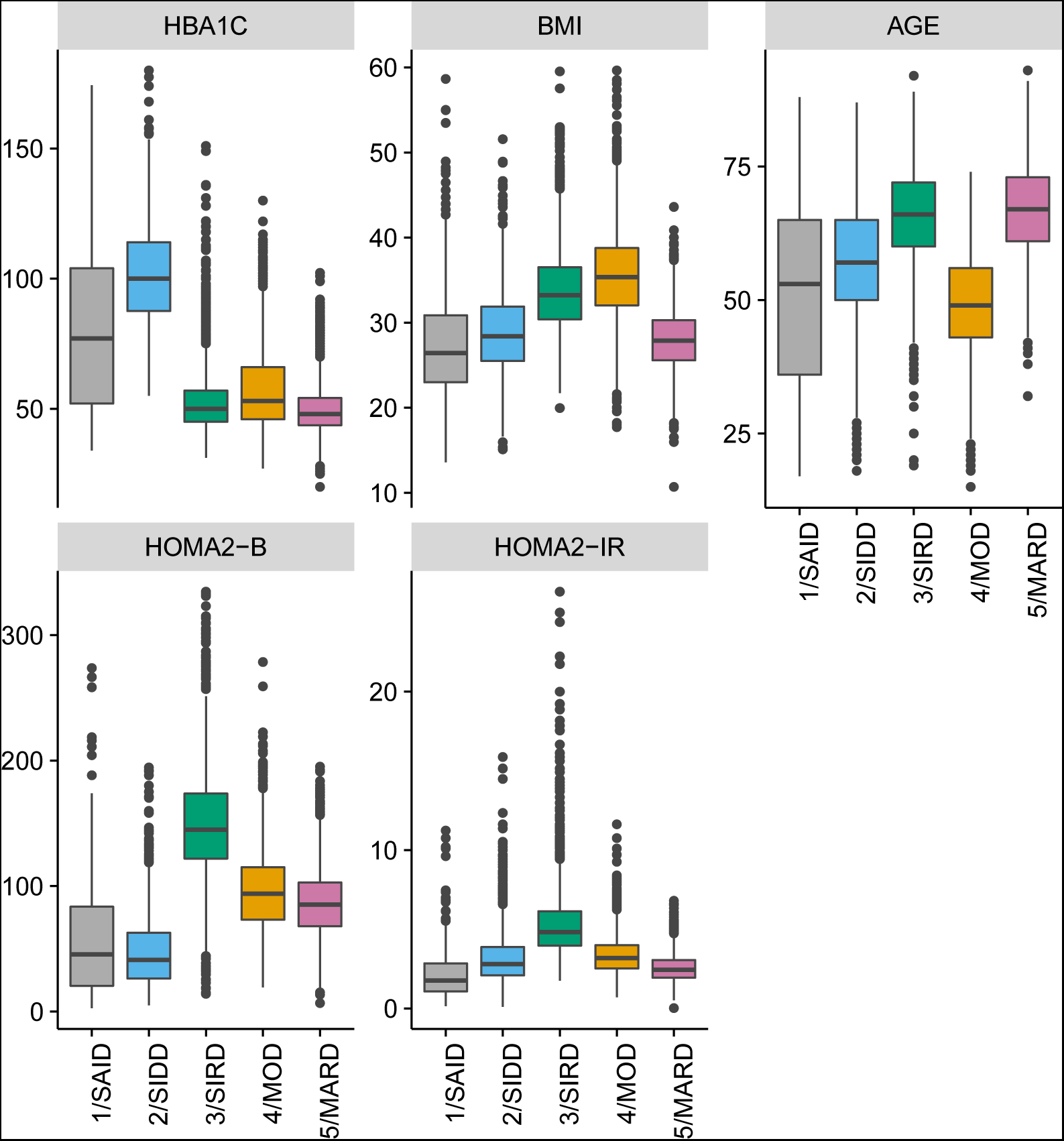
Cluster characteristics in ANDIS. Distributions of HbA1c (mmol/mol) at diagnosis, and BMI (kg/m2), age (years), HOMA2-B (%) and HOMA2-IR at registration in ANDIS for each cluster. K-means clustering was performed separately for men and women, pooled data are shown here (cluster 2-5).

Cluster 1, including 6·4% of the clustered patients (referred to as SAID, Severe Autoimmune Diabetes) was characterized by early onset, relatively low BMI, poor metabolic control, insulin deficiency, and presence of GADA (Table S2). Cluster 2 (SIDD, Severe Insulin Deficient Diabetes) encompassing 17·5% of patients was GADA negative but otherwise similar to SAID: low age at onset, relatively low BMI, low insulin secretion (low HOMA2-B) and poor metabolic control. Cluster 3 (SIRD, Severe Insulin Resistant Diabetes; 15·3%) was characterized by insulin resistance (high HOMA2-IR) and high BMI. Cluster 4 was also characterized by obesity but not by insulin resistance (MOD, Mild Obese Diabetes; 21·6%). Patients in cluster 5 were older (MARD, Mild Age-Related Diabetes; 39·1%) but showed, as cluster 4, only modest metabolic derangements.

We used three independent cohorts to replicate the clustering: the Scania Diabetes Registry (SDR, N=1,466), ANDIU (All New Diabetics in Uppsala, N=844) and DIREVA (Diabetes Registry Vaasa, N=3,485). In SDR, the optimal number of clusters was also estimated to be 5 and k-means (k=4) and TwoStep clustering yielded very similar results (92·4% clustered identically). Patient distributions and cluster characteristics were similar to ANDIS (Figure 1C, S3A and B). Jaccard bootstrap means were >0·8 for all clusters. K-means clustering in ANDIU also replicated the results from ANDIS (Figure 1D, S3D). In the DIREVA cohort we tested whether clustering would give similar results in patients with longer diabetes duration as newly-diagnosed diabetes. Encouragingly, the results were comparable in recently diagnosed patients (diabetes duration <2 years, N=878) and patients with longer duration (mean 10·15±10·34; N=2607; Figure 1E and F, S4 A and B).

To be clinically useful patients would need to be assigned to clusters without *de novo* clustering of a full cohort. Therefore, we assigned patients in replication cohorts to clusters based on which cluster they were most similar to, calculated as their Euclidian distance from the nearest cluster center derived from ANDIS coordinates and found similar distributions (Figure S3 C and E, Figure S4 B and D).

To evaluate the clinical utility of the clustering we compared disease progression, treatment and development of diabetic complications between clusters in ANDIS. SAID and SIDD had markedly higher HbA1c at diagnosis compared to other clusters, a difference persisting throughout the follow-up period (Figure 3A). Ketoacidosis at diagnosis was most frequent in SAID (30·5%) and SIDD (25·1%), compared to others (<5%, Figure S5). HbA1c was the strongest predictor of ketoacidosis at diagnosis (OR 2·73[2·46-3·03], p=2·0x10^−82^, per 1SD change, Table S4). SIRD had the highest prevalence of non-alcoholic fatty liver disease (NAFLD, Figure S6). Zinc transporter 8A antibodies were primarily seen in SAID (27·3% positive compared to <1·5% in other clusters; Figure S7).

**Figure 3.**
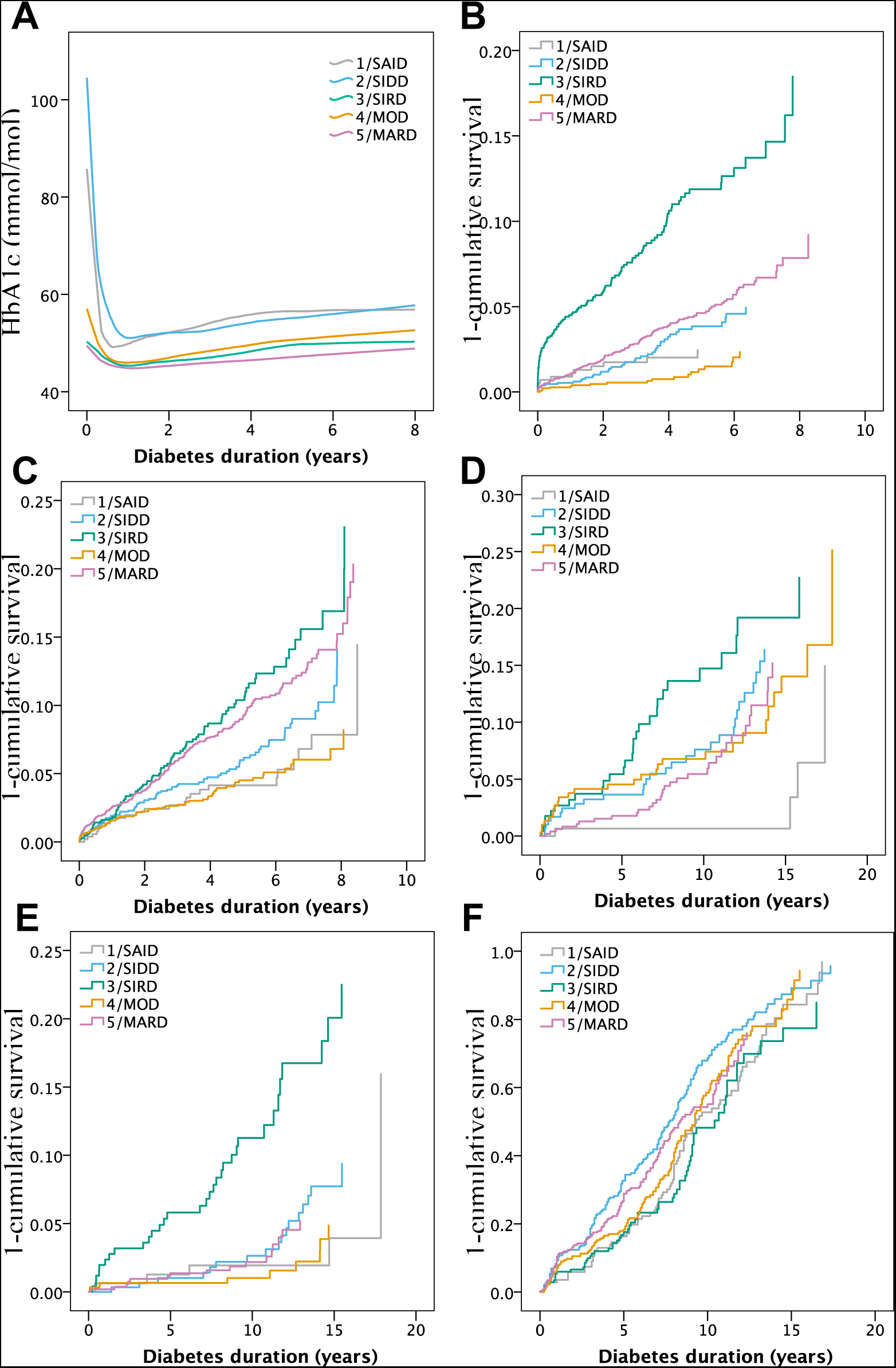
Progression of disease over time by cluster. Figure 3 shows mean HbA1c over time by loess regression (A), time to CKD at least stage 3B (B) and coronary events (C) in ANDIS; Macroalbuminuria (D), ESRD (E) and mild non-proliferative to proliferative diabetic retinopathy (F), in the SDR cohort. Kidney function was not tested at diagnosis and therefore set to the first screening date. Thus it is not known how many were already affected at diagnosis.

At registration, insulin had been prescribed to 41·9% of patients in SAID and to 29·1% in SIDD but to < 4% of patients in clusters 3-5 (Table S2, Figure S8). Time to insulin was shortest in SAID (HR 17·05[14·34-20·28] compared to MARD, Figure 4A, Table S5), followed by SIDD (HR 9·23[7·88-10·81]). The proportion of patients on metformin was highest in SIDD and lowest in SAID (Figure S8, 4B), but also surprisingly low in SIRD which should benefit most from metformin, demonstrating that traditional classification is unable to tailor treatment to the underlying pathogenic defects. Metformin is contra-indicated in patients with impaired kidney function and sometimes discontinued because of adverse gastrointestinal reactions. However, these factors had obviously not much influence on the use of metformin at this early stage of the disease, as the same distributions of patients were observed for initial treatment, and exclusion of patients with reduced kidney function (eGFR<60 ml/min) at their first check-up, had no major effect on the proportions of patients on metformin (Figure S9). Time to a second oral diabetes treatment was also shortest in SIDD (Figure 4C, Table S5). Time to reaching the treatment goal (HbA1c <52 mmol/mol) was longest for SIDD (Figure 4D).

**Figure 4.**
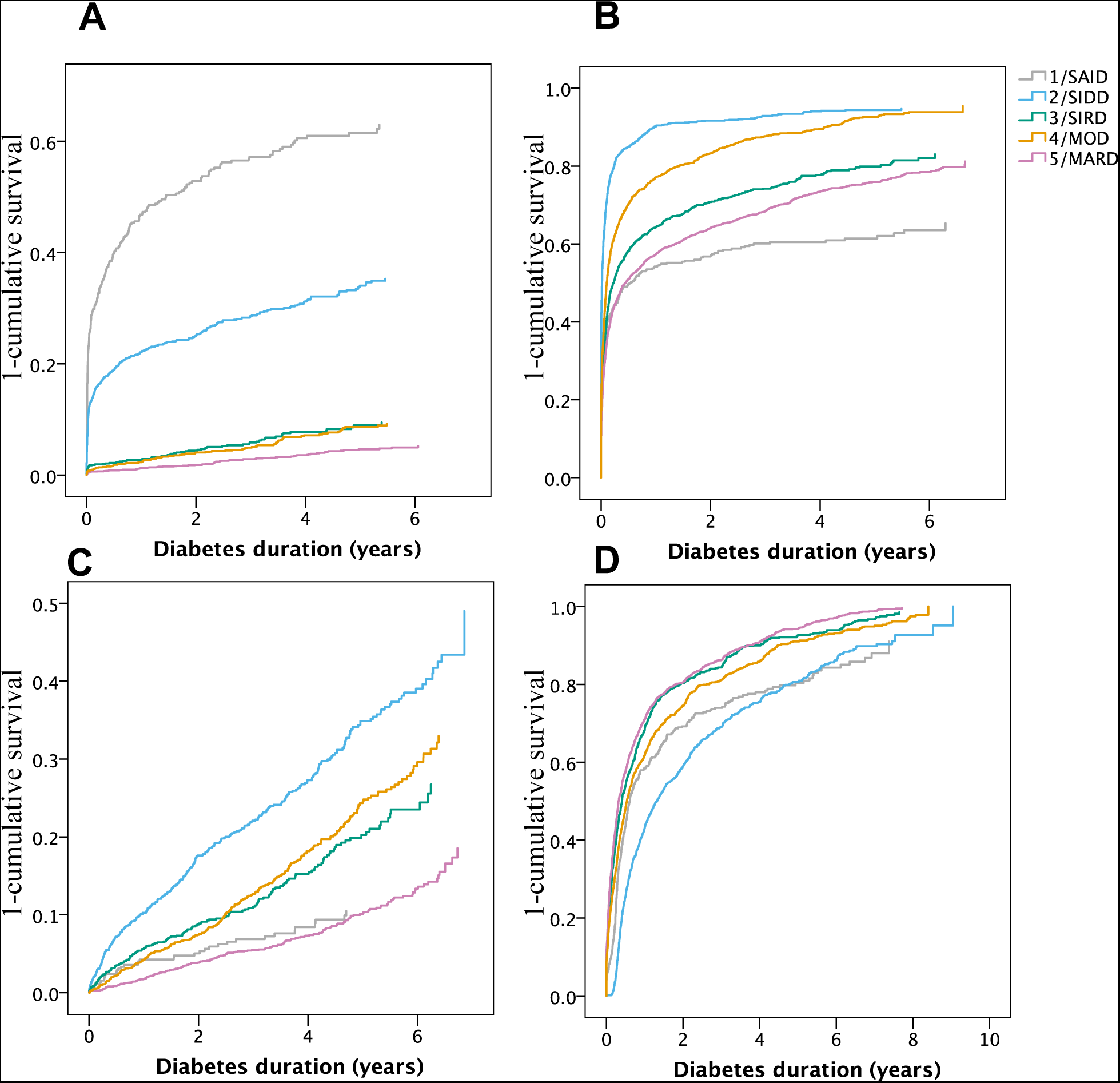
Antidiabetic therapy in ANDIS during follow-up. Cox regressions of time to treatment with insulin (A), metformin (B), oral medication other than metformin (C) or (D) reaching treatment goal (HbA1c <52mmol/mol). Cluster 1/SAID had the shortest time to insulin. Cluster 2/SIDD had a shorter time to insulin, metformin and any other oral medication than clusters 3 to 5. Despite this, cluster 2/SIDD reached the treatment goal significantly later than other clusters. For statistics see table S5.

In ANDIS, SIRD had the highest risk of developing chronic kidney disease (CKD) during follow-up of 3·9±2·3 years (Table S6). For CKD stage 3A (eGFR<60 ml/min) the age and sex adjusted risk was more than 2-fold higher (HR 2·41[2·08-2·79], p=1·4x10^−31^, Figure S10A) and for stage 3B (eGFR<45 ml/min) >3-fold higher compared with the reference cluster MARD (HR 3·34[2·59-4·30], p=8·3x10-^21^, Figure 3B). SIRD also showed higher risk of diabetic kidney disease defined as persistent macroalbuminuria, Figure S10B, HR 2·28[1·6-3·23], p=3·0x10-^6^). Also in the SDR cohort (follow-up 11·0±4·4 years), SIRD had the highest risk of CKD (Table S8), and macroalbuminuria (HR 2·36[1·54-3·61], p=7·7x10-^5^, Figure 3D). Strikingly, SIRD patients had five times higher risk of ESRD than MARD (HR 5·04[2·76-9·23], p=1·5x10-^7^, Figure 3E). The increased prevalence of kidney disease in SIRD was also confirmed in the DIREVA cohort (Figure S13).

Early signs of diabetic retinopathy (mean duration 135 days) were more common in SIDD than in other clusters (OR 1 6[1 3-1 9], p=9·7x10-^7^ compared to MARD; Figure S11A). The higher prevalence of retinopathy in SIDD was replicated in the ANDIU (Figure S1 1B) and SDR cohorts (HR 1·49[1·17-1·89], p=0·001; Figure 3F, Table S9).

Although risk of coronary events and stroke was lowest in the SAID, SIDD and MOD clusters (Figure 3C, Table S7) the differences became nominal after adjusting for age. No significant differences in age-adjusted risk of coronary events or stroke were seen between clusters in SDR (Figure S10, Table S10).

Finally, we analyzed genetic loci previously shown to be associated with T2D and related traits^17^ to see if we could obtain genetic support for the observed differences between the clusters (Table 1). Each cluster was compared to a non-diabetic cohort (MDC-CVA) from the same geographical region.^9^ Notably, no genetic variant was associated (p<0·01) with all clusters (Table S11). Strikingly, the strongest T2D-associated variant in the *TCF7L2* (rs7903146) gene^18^ was associated with SIDD, MOD and MARD, but not with SIRD (Table 1). The difference between cluster odds ratios was significant even after Bonferroni correction for the 77 tests performed. A variant in the *IGF2BP2* (rs4402960) gene was associated with SIDD and MARD but not with SIRD or MOD. The variant rs10401969 in the *TM6SF2* gene previously associated with NAFLD^19^ was most strongly associated with SIRD but not with MOD suggesting that SIRD is characterized by unhealthy (metabolic syndrome) and MOD by more healthy obesity.

## Discussion

Taken together, this study demonstrates that this new clustering is superior in terms of prediction of disease progression, particularly development of diabetic complications compared to the classical diabetes classification. Importantly, this prediction can be made already at diagnosis. In contrast to previous attempts to dissect the heterogeneity of diabetes ^20^ we have used variables reflecting key aspects of the diabetic disease that are monitored in patients. Thus, this clustering can easily be applied to both existing diabetes cohorts (e.g. from drug trials) and patients in the diabetes clinic. A web-tool to assign patients to specific clusters, provided above variables have been measured, is under development.

While SAID overlapped with T1D and LADA, SIDD and SIRD represent two novel severe forms of diabetes previously masked within T2D. It would be reasonable to target intensified treatment resources to these clusters to prevent diabetic complications. SIRD had a markedly increased risk of kidney complications, reinforcing the association between insulin resistance and kidney disease.^21^ Insulin resistance has been associated with higher salt sensitivity, glomerular hypertension, hyperfiltration, and declining renal function, all hallmarks of diabetic kidney disease (DKD).^22^ The increased incidence of DKD in this study was seen in spite of relatively low HbA1c, suggesting that glucose-lowering therapy is not the ultimate way of preventing DKD. In support of this, mice with podocyte-specific knockout of the insulin receptor, mimicking the reduced insulin signaling seen in insulin resistant individuals, developed DKD even during normoglycemic conditions.^23^ Although differences were not as pronounced as seen for DKD, insulin deficiency and/or hyperglycemia seem to be important triggers of retinopathy with the highest prevalence observed in SIDD.

We cannot at this stage claim that the new clusters represent different etiologies of diabetes, nor that this represents the optimal classification of diabetes subtypes. The fact that clustering gave similar results in newly diagnosed patients and patients with longer diabetes duration, and that the key variable C-peptide remained relatively stable over time (Figure S13), suggests that the clusters are stable and at least partially mechanistically distinct rather than representing different stages of the same disease. The differences in genetic associations also support this view. Notably, hepatic insulin resistance seems to be a feature of NAFLD, as the NAFLD-associated SNP in the *TM6SF2* gene was associated with SIRD but not with MOD. However, it still needs to be shown in prospective studies whether patients (especially from the periphery of clusters) can move between clusters and the exact overlap of weaker association signals will need to be investigated in larger cohorts. It might also be possible to refine the stratification further by including additional cluster variables e.g. biomarkers, genotypes or genetic risk scores.

Taken together, the current data demonstrate that the combined information from a few variables central to the development of diabetes is superior to the measurement of only one metabolite, glucose. By combining this information from diagnosis with information in the health care system this study provides a first step towards a more precise, clinically useful, stratification, representing an important step towards precision medicine in diabetes. This clustering also opens up for randomized trials targeting insulin secretion in SIDD and insulin resistance in SIRD.

## Acknowledgements

We thank all the patients and the health care providers across Scania and Ostrobothnia for their support and their willingness to participate. We would also like to thank Johan Hultman, Jasmina Kravic, Maria Fälemark, Christina Rosborn, Gabriella Gremsperger, Maria Sterner, Malin Neptin, Lisa Sundman, Paula Kokko, Carin Gustavsson and Ulrika Blom-Nilsson for excellent technical and administrative support. Finally we would like to thank Rita Jedlert and Region Skåne (Scania County) as well as the ANDIS steering committee for their support.

## Author contributons

EA, PS, PV, TT, AHR and LG contributed with the conception of the work. EA, PS, AK, MM, MD, AC, PV, YW, NS, PS, HM, EL, OM, OH, UM, ÅL, KL, TF, TT, AHR and LG contributed to the data collection. EA, PS, MM, RP, DMA and PA contributed to the data analysis. EA, PS and LG drafted the article. All authors contributed in the interpretation of data and critical revision of the article. All authors gave final approval of the version to be published.

---

**Figure S1.**
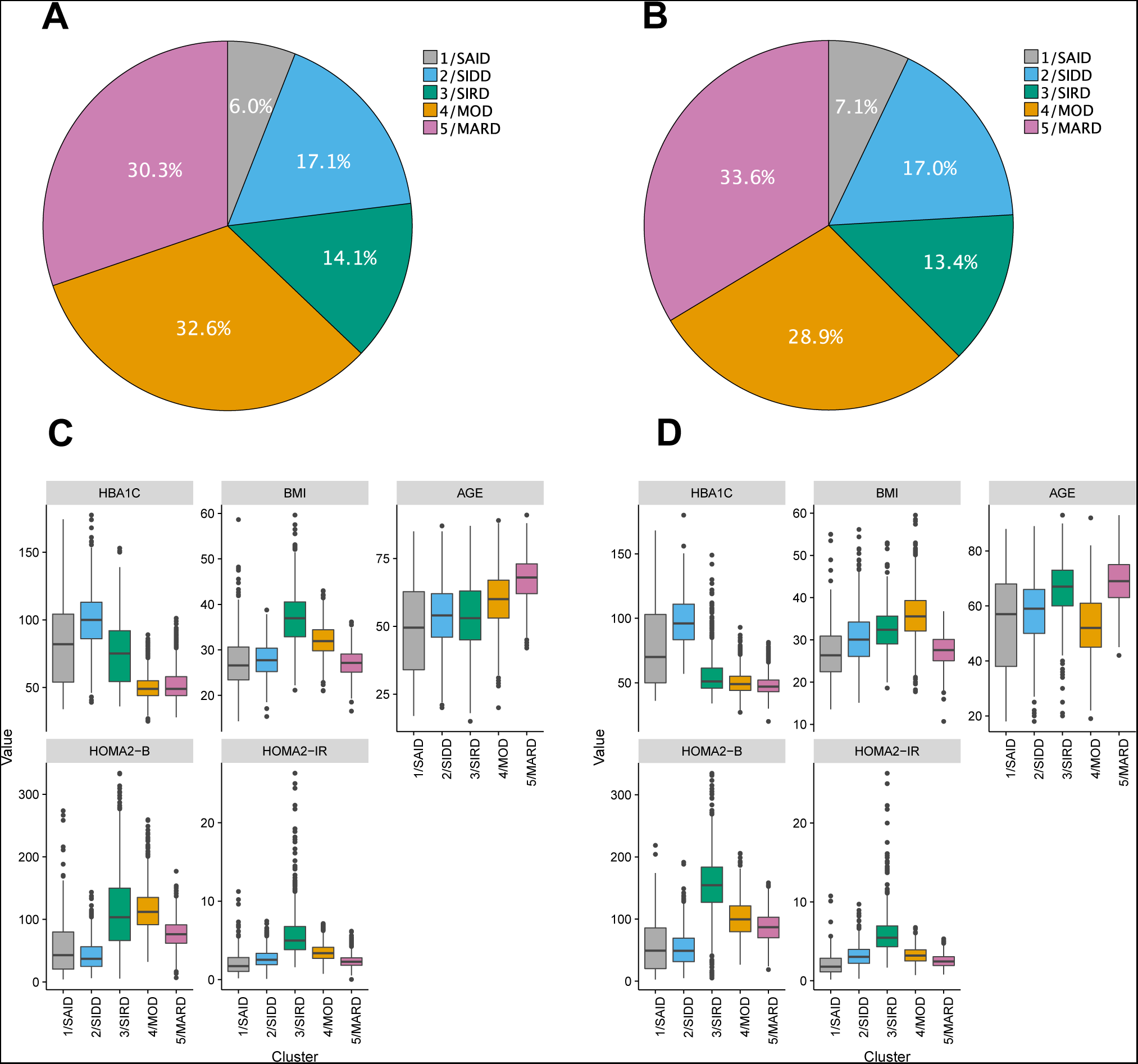
Results of sex-specific TwoStep clustering in ANDIS. The distribution of patients based on TwoStep cluster analysis in the ANDIS (All New Diabetes in Scania) cohort was similar in (A) men and (B) women. The different clusters in men (C) and women (D) had similar distribution of the variables used for clustering, i.e. of HAa1c (mmol/mol) at diagnosis; BMI (kg/m2), age (years), HOMA2-B (%) and HOMA2-IR at registration, except for some differences in clusters 3 and 4 (women in cluster 4 were older and less obese compared to men in cluster 3).

**Figure S2.**
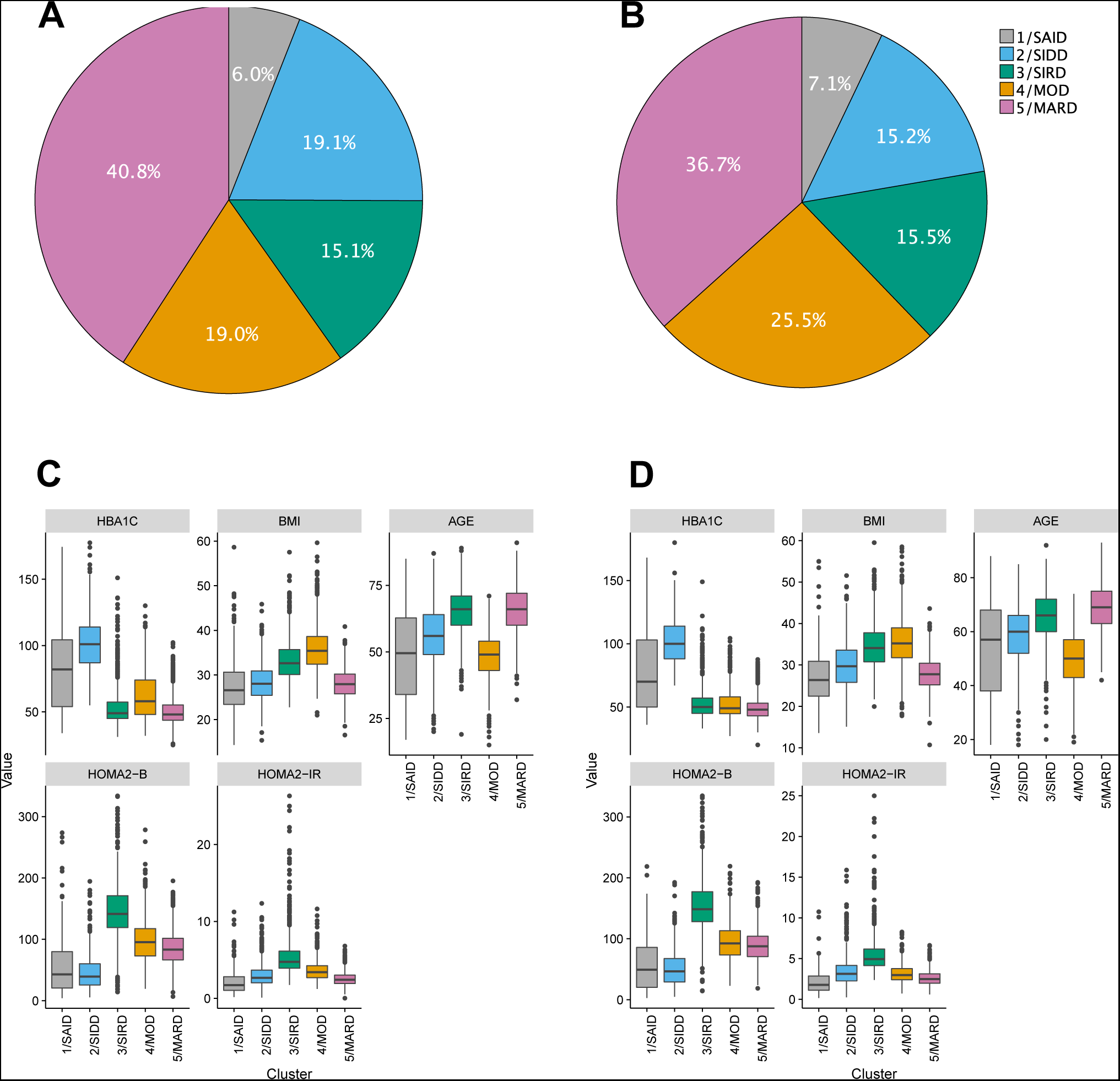
Results of sex-specific k-means clustering in ANDIS. The distribution of patients based on k-means cluster analysis in ANDIS was similar in men (A) and women (B). The different clusters in men (C) and women (D) had similar distribution of the variables used for clustering, i.e. of HAa1c (mmol/mol) at diagnosis; BMI (kg/m2), age (years), HOMA2-B (%) and HOMA2-IR at registration.

**Figure S3.**
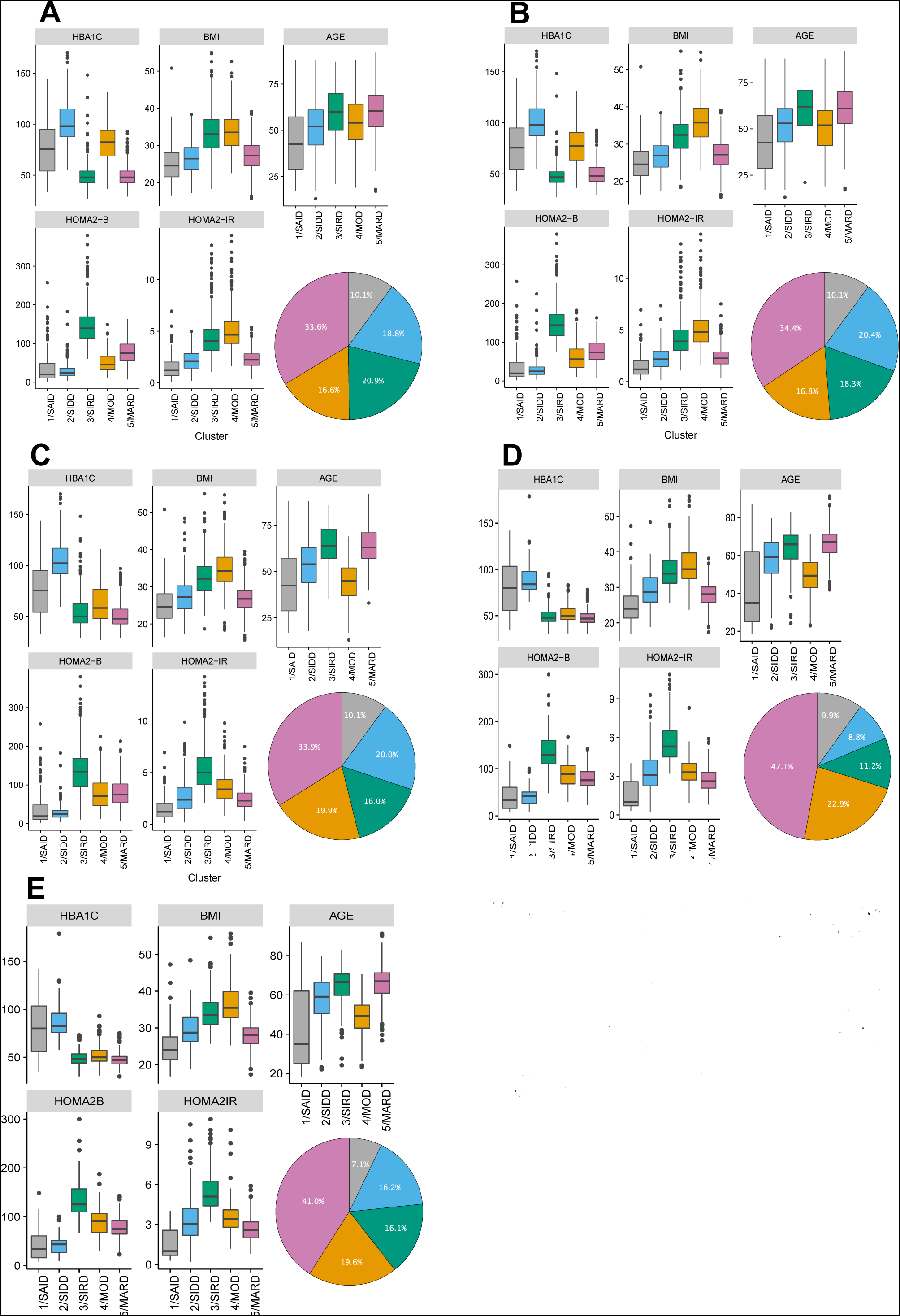
Replication of clustering in two independent Swedish cohorts, SDR and ANDIU. Cluster distributions and characteristics in the Scania Diabetes Registry (SDR) using TwoStep (A), k-means (k=4) clustering (B) and cluster assignment based on ANDIS cluster coordinates (C). Cluster distributions and characteristics in ANDIU using k-means (k=4) clustering (D) and cluster assignment D based on ANDIS cluster coordinates (E).

**Figure S4.**
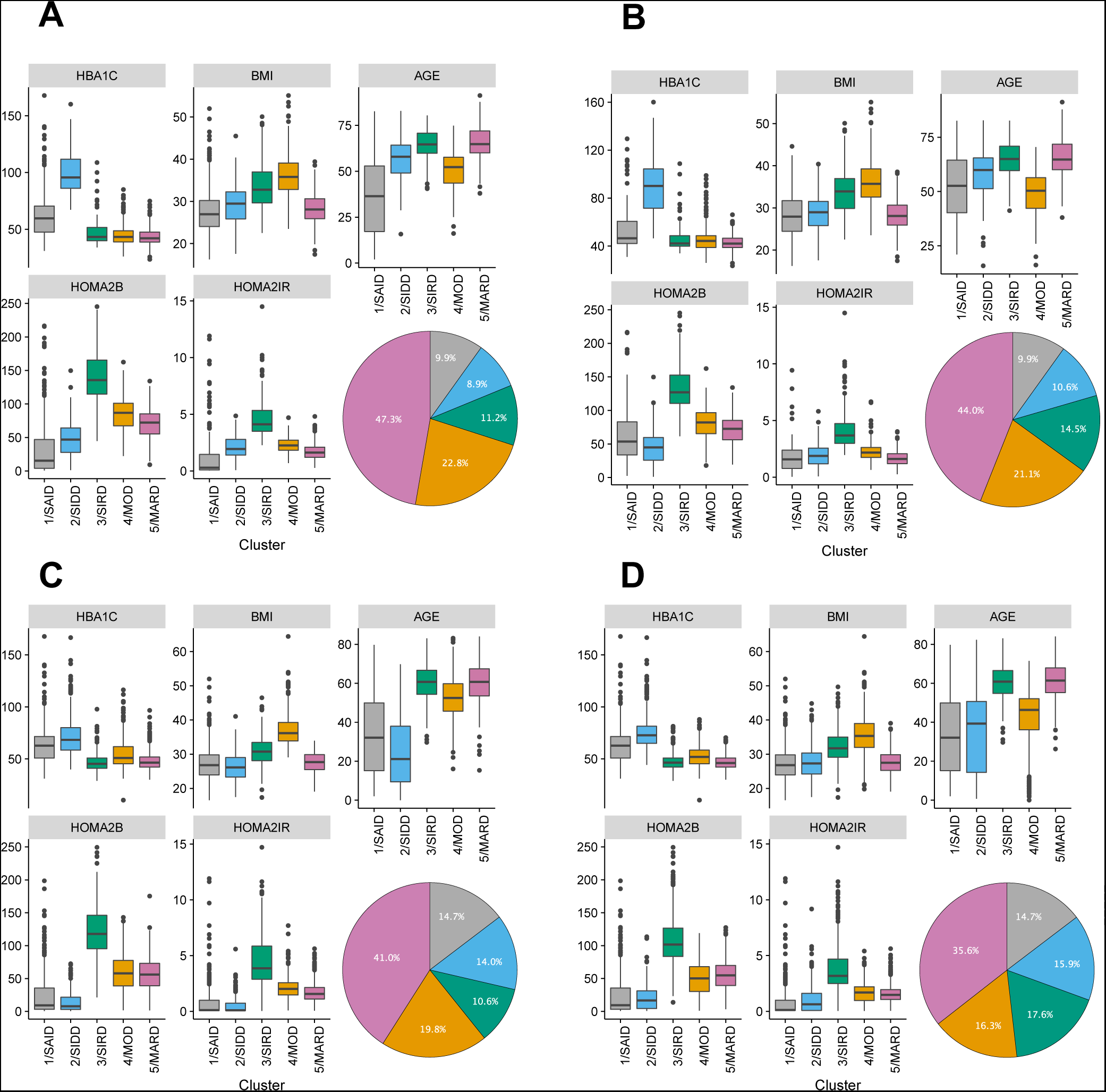
Clustering in the Finnish DIREVA cohort comparing patients with newly diagnosed diabetes and longer duration. Patient distribution in DIREVA in newly diagnosed (diabetes duration at sampling less than 2 years) using de novo k-means clustering (A), and cluster assignment based on ANDIS cluster coordinates (B). Patient distribution in patients with longer duration at sampling (mean 10·15±10·34 years) using *de novo* k-means clustering (C) and cluster assignment based on ANDIS cluster coordinates (D).

**Figure S5.**
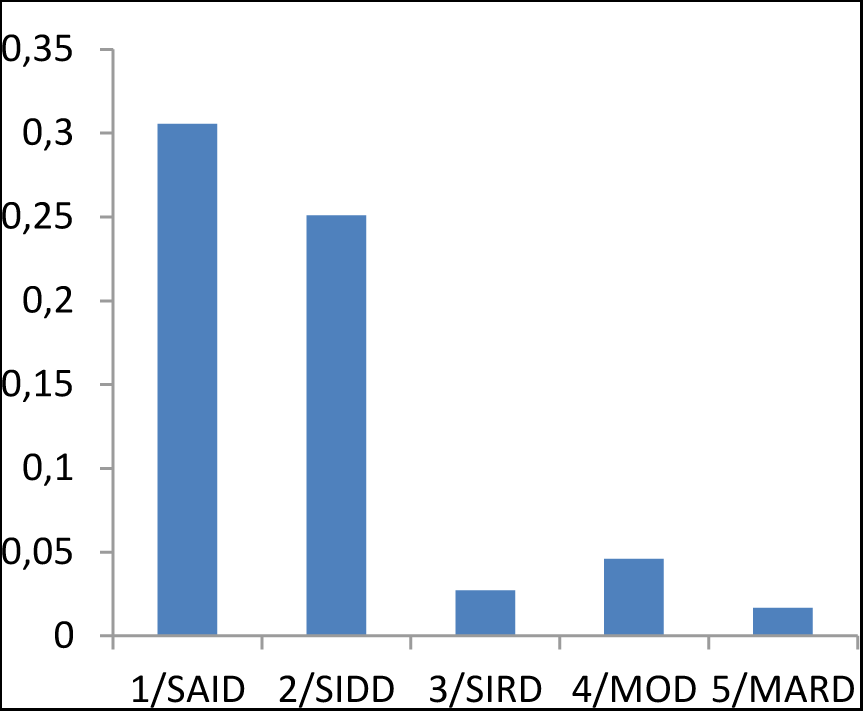
Prevalence of ketoacidosis at diagnosis in ANDIS. Ketoacidosis at diagnosis was most common in clusters 1/SAID (30·5%) and 2/SIDD (25·1%), but rare in the others (<5%).

**Figure S6.**
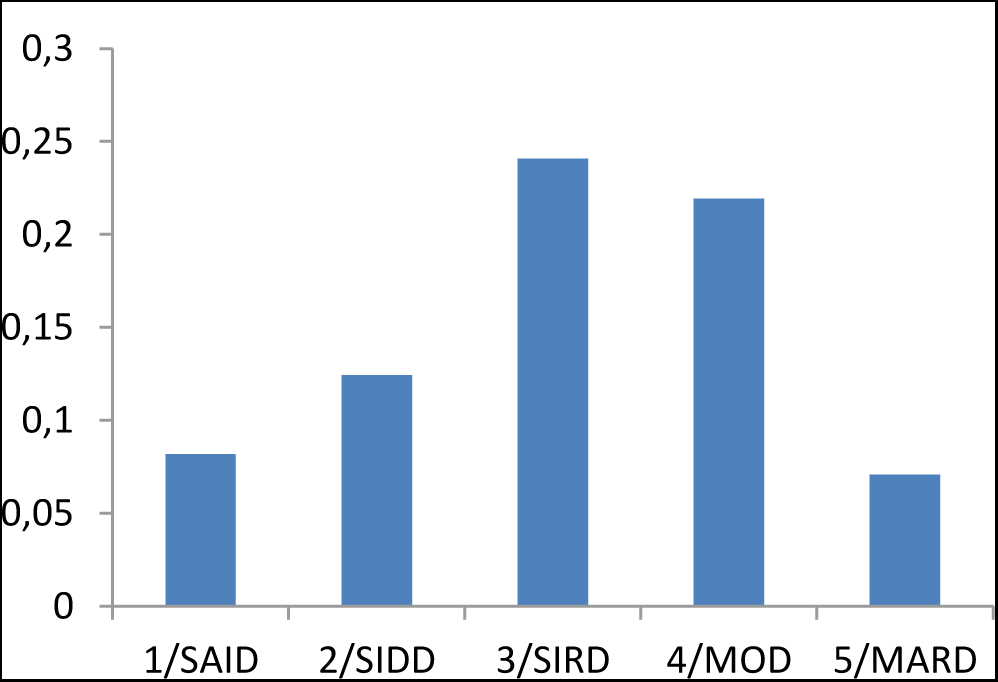
Prevalence of non-alcoholic fatty liver disease (NAFLD) in ANDIS estimated from ALT measurements. A total of 11999 patients had at least one P-ALT measurement in the database of the Clinical Chemistry unit. Of them, 3739 had at least two readings exceeding the upper reference values for the assays used (>1·1-1·2 μkat/L for men and >0·7-0·85 μkat/L for women). Cluster 3/SIRD had the highest prevalence of NAFLD, defined as two pathological ALT measurements and BMI>28 (OR 3·96[3·27-4·78], p=5·8x10^−46^ compared to MARD and OR 1·56[1·24-1·95], p=1·4x10^−4^ compared to MOD after adjustment for sex and age).

**Figure S7.**
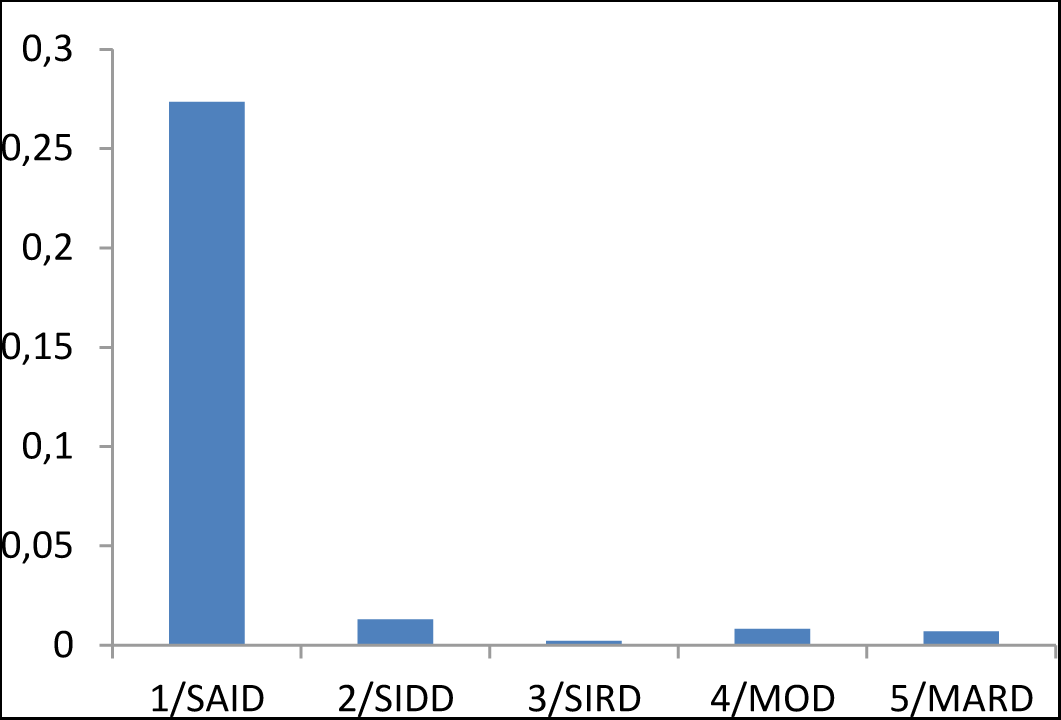
Auto-antibodies directed against zinc transporter 8A (ZnT8A). ZnT8A auto-antibody positivity was mainly observed in patients from cluster 1/SAID (OR 52·74[26·89-103·45], p=8·6x10^−31^)

**Figure S8.**
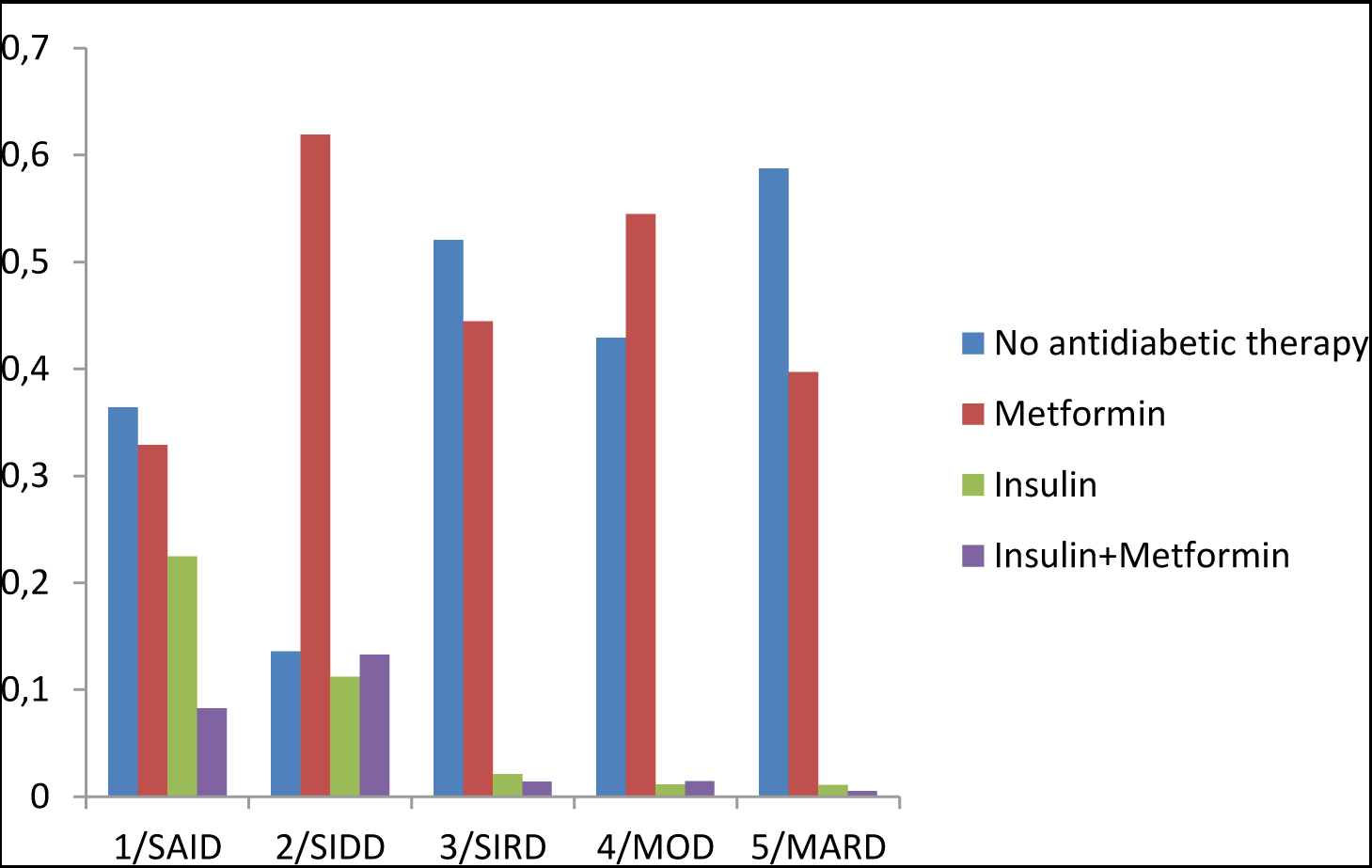
Antidiabetic medication (%) at time of registration in ANDIS. Figure S8 shows the frequency of use of metformin and/or insulin therapy at registration in ANDIS (at the time of measurement of plasma glucose and C-peptide) stratified by clustering. More patients in cluster 2/SIDD had been prescribed insulin and/or metformin than in clusters 3-5, reflecting the higher HbA1c at diagnosis.

**Figure S9.**
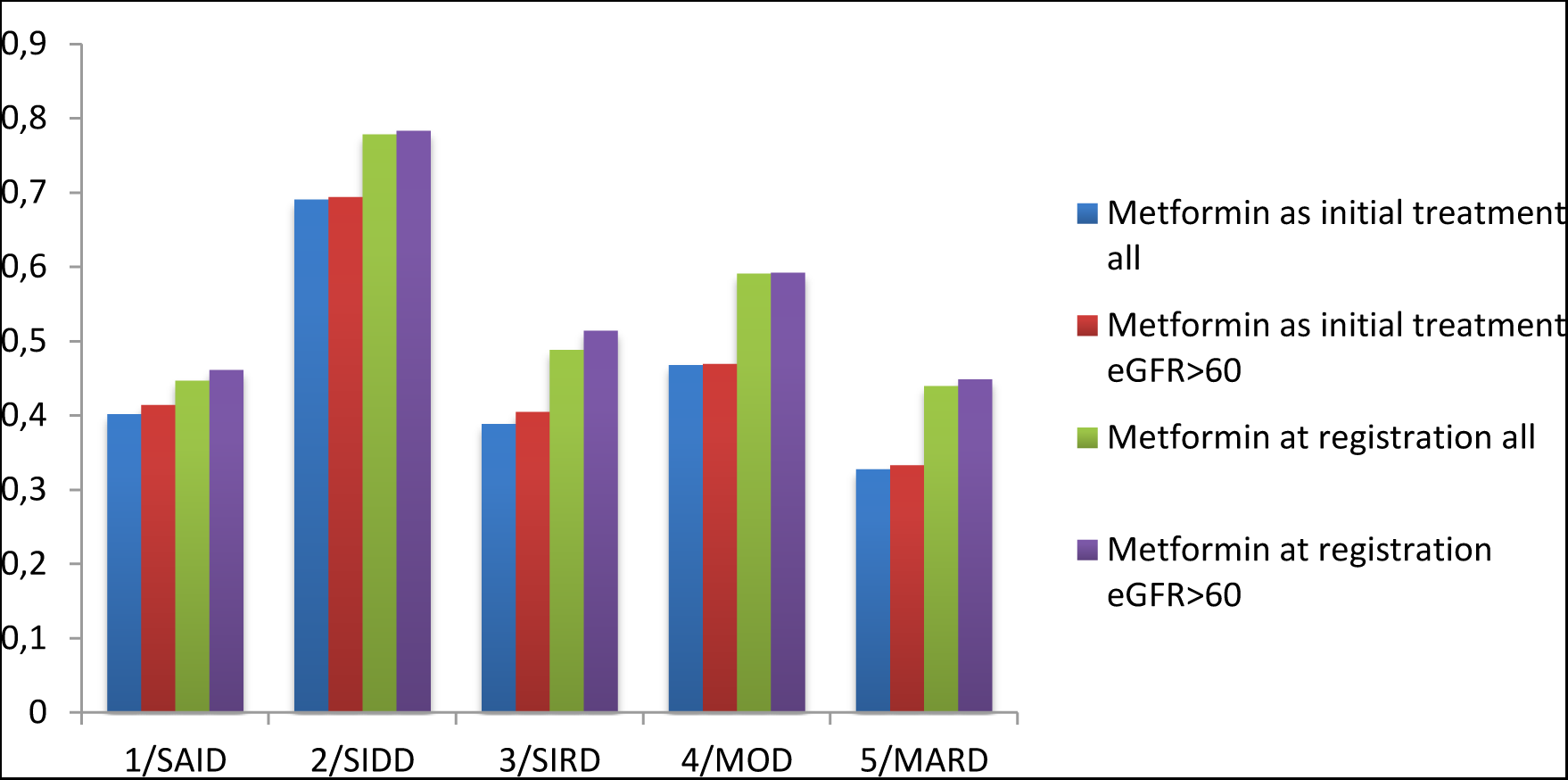
Metformin treatment (%) by cluster in ANDIS. The percentage of patients in each cluster prescribed metformin as their first treatment after diagnosis (initial treatment), and percent of renally sufficient (eGFR>60 mL/min/1·73m^2^and) patients on metformin at registration. This shows that the difference between clusters is not a result of discontinuation of metformin due to adverse effects or contra-indication of metformin in patients with kidney disease.

**Figure S10.**
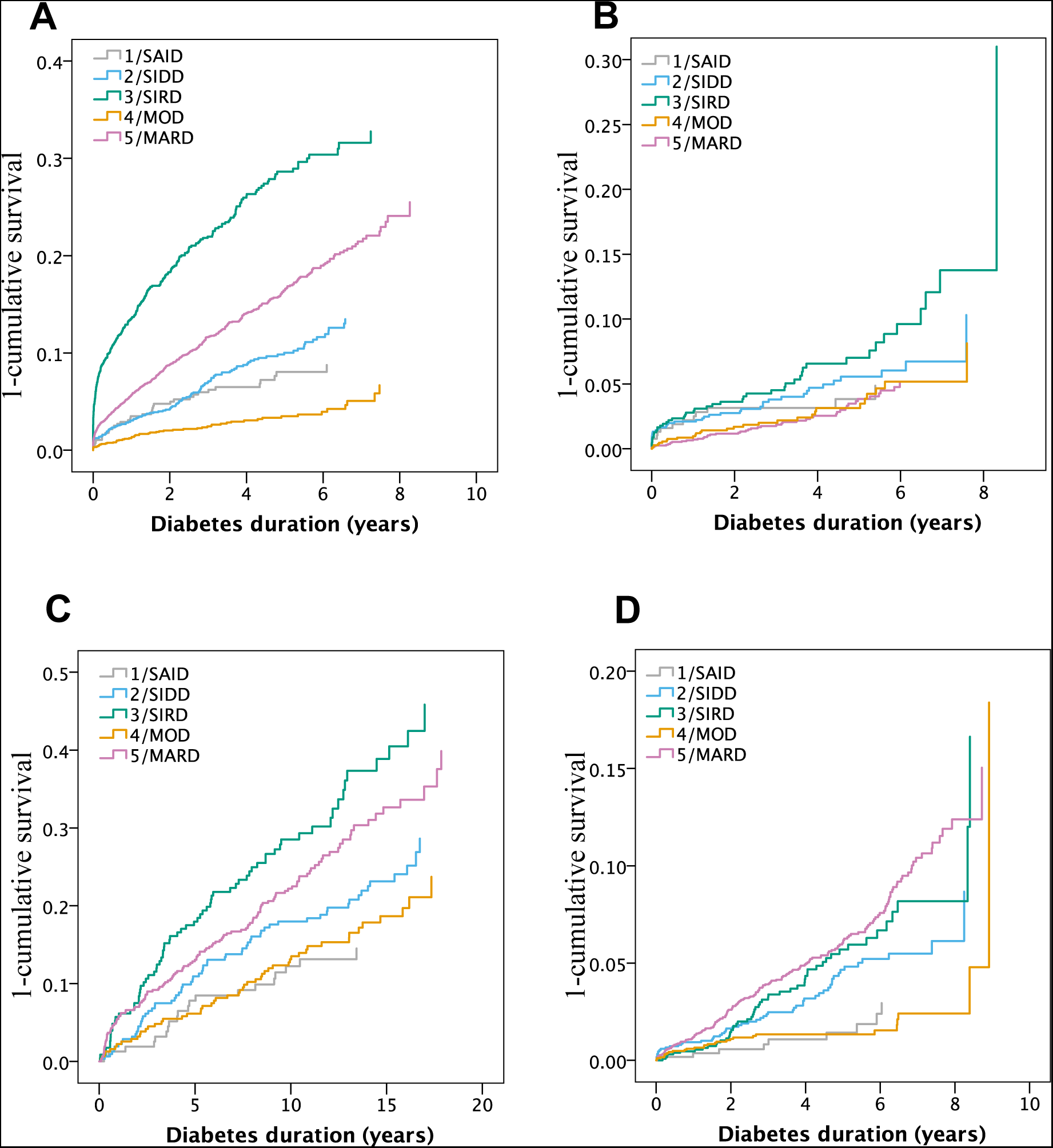
Risk of diabetic complications by cluster. Cox regressions of diabetic complications for (A) CKD stage 3A (eGFR <60 ml/min) and macroalbuminuria (B) in ANDIS, coronary events (C) and stroke (D) in SDR. For statistics see Table S6 and S10.

**Figure S11.**
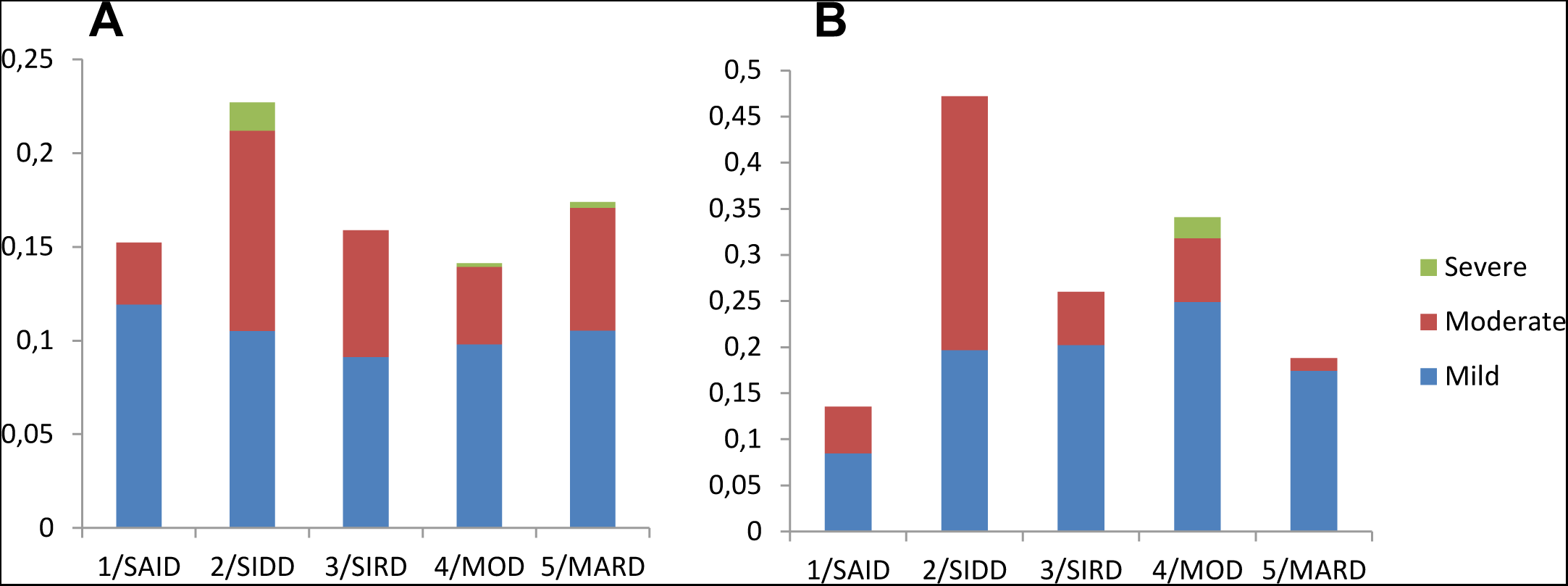
Diabetic retinopathy (DR) at diagnosis. Prevalence of different stages of diabetic retinopathy in (A) ANDIS and (B) ANDIU. In ANDIS DR risk was significantly higher in SIDD than in the reference cluster MARD (OR 1·6[1·3-1·9], p=9·7x10^−7^). In ANDIU DR risk was elevated in both SIDD (OR 4·6[3·0-7·0], p=4·1x10^−13^) and MOD compared to MARD (OR 2·2[1·4-3·3], p=2·8x10^−4^).

**Figure S12.**
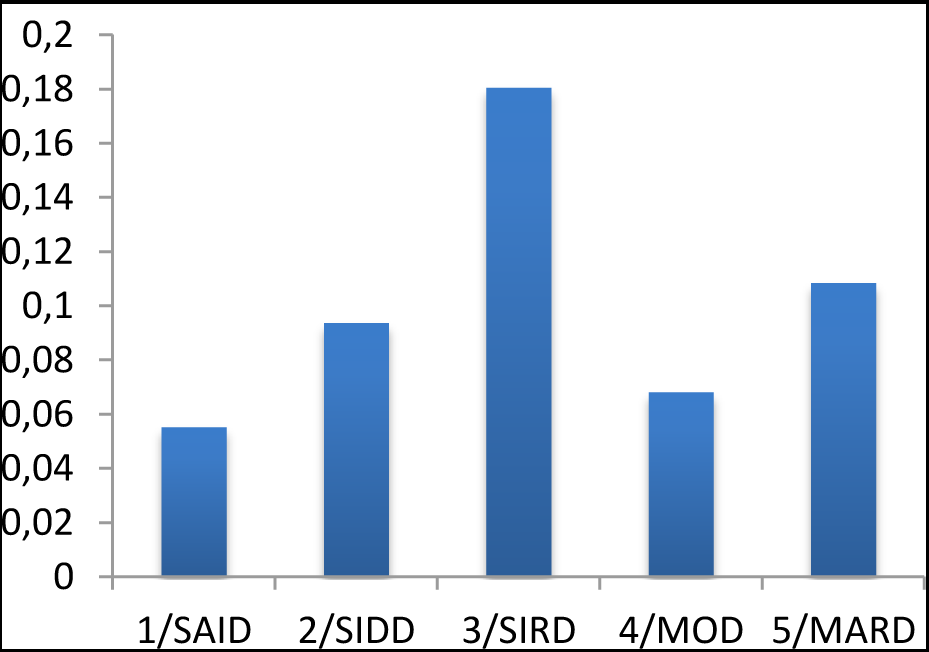
Prevalence of chronic kidney disease in DIREVA. CKD (eGFR<60 ml/min) in DIREVA by clusters assigned based on ANDIS cluster coordinates. SIRD had increased risk of CKD (OR 2·02[1·38-2·96], p=3·4x10^−4^) after adjustment for age, sex and duration of diabetes.

**Figure S13.**
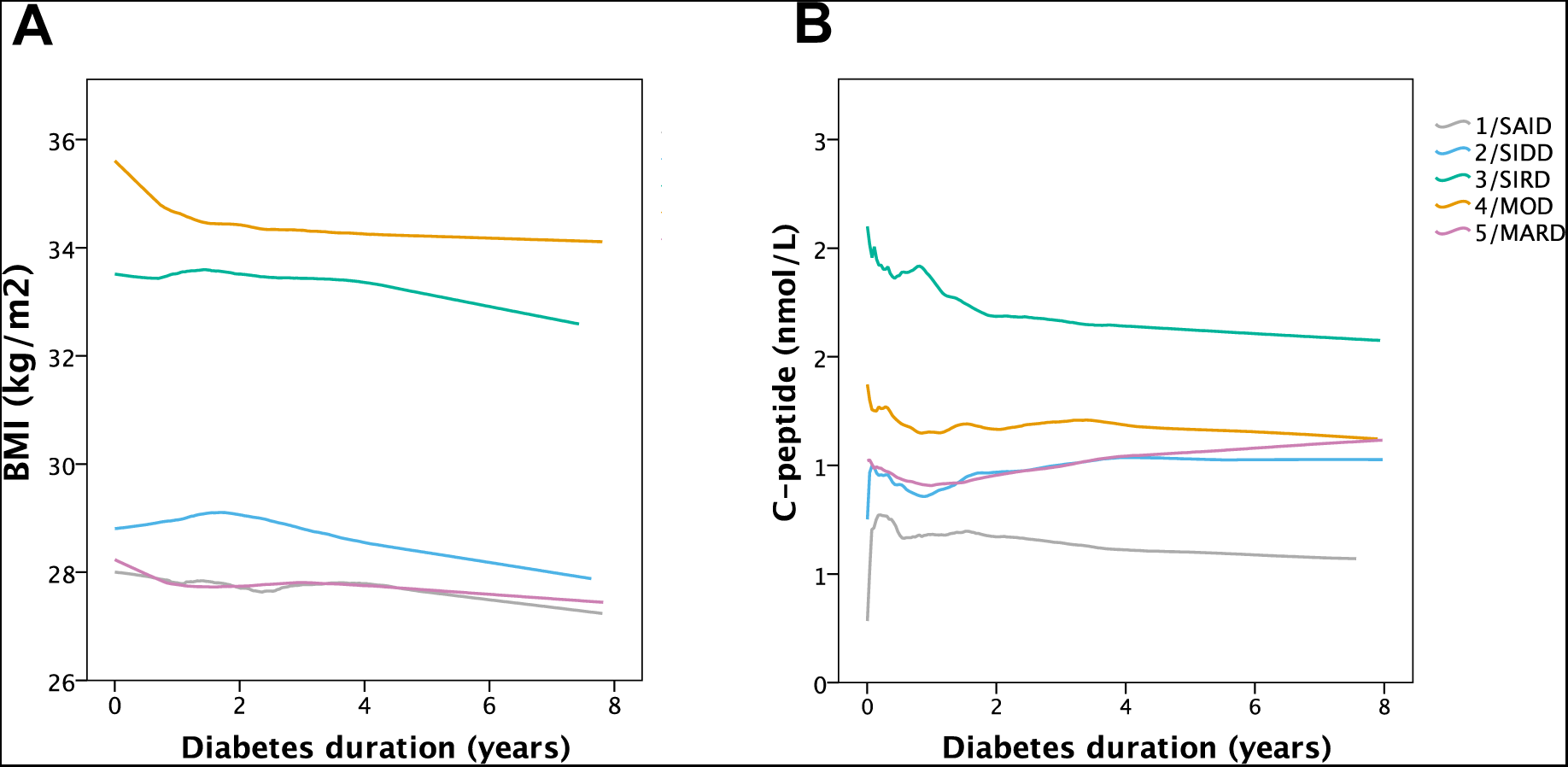
Change in cluster variables over time in ANDIS. Figure S13 shows change in BMI(A) and C-peptide (B) during follow-up by cluster.

**Table S1.** Patient characteristics in ANDIS using the traditional classification*

|  | **T1D** | **LADA** | **T2D** |
| --- | --- | --- | --- |
| **N** | 204 | 723 | 12112 |
| **Frequency in total cohort, %** | 1.4 | 4.9 | 82.7 |
| **Men, %** | 64.7 | 54.1 | 59.7 |
| **HBA1C at diagnosis, mmol/l** | 107.88(27.06) | 73.65(28.26) | 62.32(24.12) |
| **BMI, kg/m2** | 22.02(3.50) | 28.77(6.32) | 30.93(5.72) |
| **Age at diagnosis, years** | 34.03(13.84) | 54.86(12.67) | 60.93(12.25) |
| **HOMA2B** | 23.50(20.94) | 65.46(46.88) | 91.95(48.19) |
| **HOMA2IR** | 0.66(0.35) | 2.64(2.08) | 3.41(2.55) |
| **Insulin at registration, %** | 96.5 | 33.4 | 9.9 |
| **Metformin at registration, %** | 10.2 | 43.8 | 52.0 |
| **History of gestational diabetes, % of women** | 3.9 | 11.9 | 11.3 |
| **Non-Scandinavian origin %** | 12.5 | 16.8 | 23.2 |

\*Only patients older than 18 at registration are included

**Table S2.** Patient characteristics in ANDIS using k-means clustering.

|  | **SAID** | **SIDD** | **SIRD** | **MOD** | **MARD** |
| --- | --- | --- | --- | --- | --- |
| **N** | 577 | 1575 | 1373 | 1942 | 3513 |
| **Frequency, %** | 6.4 | 17.5 | 15.3 | 21.6 | 39.1 |
| **Men, %** | 55.1 | 64.8 | 58.8 | 52.1 | 62.0 |
| **HBA1C at diagnosis, mmol/l** | 80.03(30.84) | 101.85(19.26) | 54.07(15.46) | 57.70(16.07) | 50.08(9.85) |
| **BMI, kg/m2** | 27.45(6.44) | 28.86(4.77) | 33.85(5.24) | 35.71(5.43) | 27.94(3.44) |
| **Age at diagnosis, years** | 50.48(17.93) | 56.74(11.14) | 65.25(9.34) | 48.96(9.54) | 67.37(8.55) |
| **HOMA2B** | 56.71(44.65) | 47.64(28.93) | 150.47(47.20) | 95.03(32.45) | 86.59(26.37) |
| **HOMA2IR** | 2.16(1.56) | 3.18(1.73) | 5.54(2.74) | 3.35(1.21) | 2.55(0.84) |
| **Insulin at registration, %** | 41.9 | 29.1 | 3.7 | 3.3 | 1.6 |
| **Metformin at registration, %** | 44.7 | 77.8 | 48.8 | 59.1 | 44.0 |
| **Family history of diabetes, %** | 59 | 64 | 56 | 70 | 58 |
| **History of gestational diabetes, % of women** | 10.3 | 7.5 | 4.5 | 21.7 | 5 |
| **Non-Scandinavian origin %** | 15.4 | 26.6 | 15.1 | 32.3 | 16.8 |

**Table S3.** Cluster centres in ANDIS

|  | Cluster | HbA1C | BMI | AGE | HOMA2B | HOMA2IR |
| --- | --- | --- | --- | --- | --- | --- |
| Women | 2/SIDD | 1·8702613 | -0·2415449 | -0·1929637 | -0·97446899 | 0·056469 |
|  | 3/SIRD | -0·254848 | 0·5189057 | 0·3214557 | 1·35581907 | 1·1801933 |
|  | 4/MOD | -0·3003478 | 0·6683606 | -0·9388278 | -0·03556857 | -0·1405151 |
|  | 5/MARD | -0·4582762 | -0·5854255 | 0·5980588 | -0·14552652 | -0·4254893 |
| Men | 2/SIDD | 1·52185804 | -0·4284673 | -0·4017103 | -0·98397328 | -0·1630751 |
|  | 3/SIRD | -0·39080167 | 0·5396294 | 0·4235841 | 1·29059153 | 1·1801031 |
|  | 4/MOD | -0·06915764 | 1·0305317 | -1·0157681 | 0·15742215 | 0·1343923 |
|  | 5/MARD | -0·5367578 | -0·4776681 | 0·5031031 | -0·09004338 | -0·4233873 |

**Table S4.** Logistic regression analysis of risk factors for ketoacidosis at diagnosis in ANDIS

|  | OR(CI95%)* | P |
| --- | --- | --- |
| **HOMA2-B** | 0·66(0·56-0·77) | 1·1x10<sup>-7</sup> |
| **HOMA2-IR** | 0·95(0·81-1·12) | 0·55 |
| **Age** | 0·65(0·59-0·72) | 1·5x10<sup>-17</sup> |
| **BMI** | 0·94(0·84-1·05) | 0·28 |
| **HbA1c** | 2·73(2·47-3·03) | 2·0x10<sup>-82</sup> |

\*OR are for 1 SD change in variable.

**Table S5.** Cox regression analysis comparing antidiabetic treatments between clusters in ANDIS.

|  | Events | Censored | % events | HR(CI95%) | P |
| --- | --- | --- | --- | --- | --- |
| **Sustained insulin** |  |  |  |  |  |
| 1/SAID | 260 | 194 | 57.3 | 26.87(21.17-34.11) | 4.3x10<sup>-161</sup> |
| 2/SIDD | 381 | 908 | 29.6 | 10.97(8.73-13.77) | 2.5x10<sup>-94</sup> |
| 3/SIRD | 73 | 1121 | 6.1 | 2.03(1.49-2.76) | 7.0x10<sup>-6</sup> |
| 4/MOD | 97 | 1539 | 5.9 | 1.90(1.43-2.53) | 1.0x10<sup>-5</sup> |
| 5/MARD | 92 | 2806 | 3.2 | 1 | - |
| **Metformin** |  |  |  |  |  |
| 1/SAID | 267 | 177 | 60.1 | 0.83(0.73-0.95) | 0.005 |
| 2/SIDD | 1152 | 81 | 93.4 | 2.56(2.38-2.76) | 2.7x10<sup>-138</sup> |
| 3/SIRD | 859 | 295 | 74.4 | 1.17(1.08-1.27) | 8.5x10<sup>-5</sup> |
| 4/MOD | 1402 | 185 | 88.3 | 1.67(1.56-1.79) | 1.8x10<sup>-48</sup> |
| 5/MARD | 2007 | 826 | 70.8 | 1 | - |
| **Oral treatment other than metformin** |  |  |  |  |  |
| 1/SAID | 36 | 419 | 7.9 | 1.00(0.70-1.42) | 0.99 |
| 2/SIDD | 339 | 950 | 26.3 | 3.99(3.36-4.74) | 3.0x10<sup>-56</sup> |
| 3/SIRD | 162 | 1034 | 13.5 | 2.10(1.71-2.58) | 1.1x10<sup>-12</sup> |
| 4/MOD | 264 | 1372 | 16.1 | 2.36 (1.97-2.83) | 2.1x10<sup>-20</sup> |
| 5/MARD | 212 | 2687 | 7.3 | 1 | - |
| **Reaching treatment goal** |  |  |  |  |  |
| 1/SAID | 353 | 130 | 73.1 | 0.60(0.53-0.67) | 9.4x10<sup>-19</sup> |
| 2/SIDD | 957 | 516 | 65.0 | 0.46(0.43-0.50) | 1.3x10<sup>-84</sup> |
| 3/SIRD | 726 | 109 | 86.9 | 0.90(0.83-0.98) | 0.016 |
| 4/MOD | 1099 | 257 | 81.0 | 0.77(0.71-0.83) | 5.0x10<sup>-12</sup> |
| 5/MARD | 1905 | 240 | 88.8 | 1 | - |

The table information from the Swedish Drug Prescription Registry of patient’s pick-up of medication from the pharmacy. Sustained insulin is defined as more than 6 months on insulin treatment. Reaching treatment goal is defined as the first time point after diagnosis the HbA1c is below 52 mmol/mol.

**Table S6.** Cox regression analysis comparing risk of kidney complications in ANDIS

|  |  | Events | Censored | Events<br>(%) | HR(CI95%)<sup>1</sup> | p<sup>1</sup> | HR(CI95%)<sup>2</sup> | p<sup>2</sup> |
| --- | --- | --- | --- | --- | --- | --- | --- | --- |
| **CKD 3A** | 1/SAID | 37 | 534 | 6.5 | 0.91(0.65-1.27) | 0.57 | 0.86(0.61-1.20) | 0.37 |
|  | 2/SIDD | 123 | 1411 | 8.0 | 1.34(1.09-1.64) | 0.006 | 1.23(1.00-1.51) | 0.06 |
|  | 3/SIRD | 298 | 1037 | 22.3 | 2.41(2.08-2.79) | 1.4x10<sup>-31</sup> | 1.56(1.34-1.82) | 6.4x10<sup>-9</sup> |
|  | 4/MOD | 55 | 1828 | 2.9 | 1.17(0.86-1.60) | 0.31 | 1.06(0.77-1.44) | 0.73 |
|  | 5/MARD | 454 | 2950 | 13.3 | 1 | - | 1 | - |
| **CKD 3B** | 1/SAID | 11 | 557 | 1.9 | 1.01(0.55-1.87) | 0.98 | 0.92(0.49-1.70) | 0.78 |
|  | 2/SIDD | 42 | 1492 | 2.7 | 1.73(1.21-2.48) | 0.003 | 1.21(0.83-1.76) | 0.32 |
|  | 3/SIRD | 118 | 1217 | 8.8 | 3.34(2.59-4.30) | 8.3x10<sup>-21</sup> | 1.86(1.44-2.41) | 3.0x10<sup>-6</sup> |
|  | 4/MOD | 19 | 1821 | 1.0 | 1.73(1.01-2.96) | 0.045 | 1.49(0.86-2.57) | 0.15 |
|  | 5/MARD | 128 | 3270 | 3.8 | 1 | - | 1 | - |
| **Macro-albuminuria** | 1/SAID | 13 | 379 | 3.3 | 1.48(0.80-2.74) | 0.21 | 2.52(1.37-4.79) | 0.005 |
|  | 2/SIDD | 41 | 970 | 4.1 | 1.80(1.19-2.74) | 0.006 | 2.71(1.74-4.20) | 9.0x10<sup>-6</sup> |
|  | 3/SIRD | 46 | 770 | 5.6 | 2.74(1.82-4.11) | 1.0x10<sup>-6</sup> | 2.45(1.62-3.71) | 2.2x10<sup>-5</sup> |
|  | 4/MOD | 29 | 1077 | 2.6 | 1.26(0.79-2.00) | 0.34 | 2.19(1.28-3.75) | 0.004 |
|  | 5/MARD | 47 | 1964 | 2.3 | 1 | - | 1 | - |
| **ESRD** | 1/SAID | 3 | 482 | 0.6 | 1.16(0.35-3.84) | 0.81 | 1.07(0.33-3.55) | 0.91 |
|  | 2/SIDD | 16 | 1438 | 1.1 | 2.58(1.36-4.91) | 0.004 | 2.68(1.42-5.07) | 0.002 |
|  | 3/SIRD | 25 | 1306 | 1.9 | 3.12(1.82-5.36) | 3.8x10<sup>-5</sup> | 2.00(1.15-3.49) | 0.015 |
|  | 4/MOD | 8 | 1313 | 0.6 | 2.37(0.96-5.83) | 0.06 | 2.04(0.82-5.05) | 0.12 |
|  | 5/MARD | 28 | 3203 | 0.9 | 1 | - | 1 | - |

<sup>1</sup> Cox regressions adjusted for sex and age at onset.
<sup>2</sup> Cox regression adjusted for sex, age at onset and first eGFR
CKD3A was defined as eGFR <60 mL/min/1.73m<sup>2</sup> and CKD3B as eGFR <45 mL/min/1.73m<sup>2</sup> for more than 90 days. End-stage renal disease (ESRD) was defined as at least one eGFR below 15 mL/min/1.73m<sup>2</sup>. Diabetic kidney disease (DKD) was defined as either ESRD or macroalbuminuria defined as at least two out of three consecutive visits with albumin excretion rate (AER) ≥200 µg/min or AER ≥300 mg/24 h or albumin-creatinine ratio (ACR) ≥25/35 mg/mmol for men/women. Median duration at first eGFR 53(IQR 1-286) days.

**Table S7.** Cox regression analysis comparing risk of cardiovascular disease between clusters in ANDIS

|  |  | Events | Censored | Events<br>(%) | HR(CI95%)<sup>1</sup> | p<sup>1</sup> | HR(CI95%)<sup>2</sup> | p<sup>2</sup> |
| --- | --- | --- | --- | --- | --- | --- | --- | --- |
| **CE** | 1/SAID | 24 | 504 | 4.5 | 0.51(0.34-0.78) | 0.002 | 0.94(0.61-1.45) | 0.77 |
|  | 2/SIDD | 82 | 1377 | 5.6 | 0.66(0.51-0.85) | 0.001 | 0.93(0.71-1.22) | 0.60 |
|  | 3/SIRD | 89 | 979 | 8.3 | 1.18(0.92-1.50) | 0.20 | 1.29(1.00-1.65) | 0.047 |
|  | 4/MOD | 66 | 1706 | 3.7 | 0.49(0.37-0.64) | 3.3x10<sup>-7</sup> | 1.15(0.80-1.65) | 0.45 |
|  | 5/MARD | 227 | 2604 | 8.0 | 1 | - | 1 | - |
| **Stroke** | 1/SAID | 9 | 560 | 1.6 | 0.25(0.13-0.49) | 4.6x10<sup>-5</sup> | 0.42(0.22-0.84) | 0.013 |
|  | 2/SIDD | 54 | 1474 | 3.5 | 0.61(0.45-0.83) | 0.001 | 0.82(0.70-1.33) | 0.82 |
|  | 3/SIRD | 53 | 1250 | 4.1 | 0.81(0.60-1.10) | 0.18 | 0.88(0.65-1.19) | 0.40 |
|  | 4/MOD | 28 | 1851 | 1.5 | 0.26(0.18-0.39) | 5.4x10<sup>-11</sup> | 0.56(0.35-0.91) | 0.019 |
|  | 5/MARD | 191 | 3184 | 5.7 | 1 | - | 1 | - |

<sup>1</sup> Cox regression adjusted for sex.
<sup>2</sup> Cox regressions adjusted for sex and age at onset.
Coronary events (CE) were defined by ICD-10 codes I21, I252, I20, I251, I253, I254, I255, I256, I257, I258, I259. Stroke was defined by ICD-10 codes I60, I61, I63 and I64.

**Table S8.** Cox regression analysis comparing risk of kidney complications between clusters in SDR

|  |  | Events | Censored | Events (%) | HR(CI95%)<sup>1</sup> | p<sup>1</sup> | HR(CI95%)<sup>2</sup> | p<sup>2</sup> |
| --- | --- | --- | --- | --- | --- | --- | --- | --- |
| **CKD 3A** | 1/SAID | 24 | 140 | 14.6 | 1.08(0.70-1.66) | 0.74 | 1.01(0.65-1.56) | 0.97 |
|  | 2/SIDD | 85 | 235 | 26.6 | 1.50(1.14-1.96) | 0.003 | 1.13(0.85-1.49) | 0.40 |
|  | 3/SIRD | 118 | 141 | 45.6 | 1.87(1.48-2.38) | 2.4x10<sup>-7</sup> | 1.39(1.09-1.76) | 0.008 |
|  | 4/MOD | 43 | 272 | 13.7 | 1.41(0.97-2.06) | 0.073 | 1.47(1.00-2.16) | 0.05 |
|  | 5/MARD | 161 | 382 | 29.7 | 1 | - | 1 | - |
| **CKD 3B** | 1/SAID | 14 | 149 | 8.6 | 1.25(0.70-2.22) | 0.45 | 1.03(0.58-1.84) | 0.92 |
|  | 2/SIDD | 51 | 269 | 15.9 | 1.75(1.21-2.52) | 0.003 | 1.24(0.85-1.81) | 0.26 |
|  | 3/SIRD | 69 | 190 | 26.6 | 2.19(1.58-3.04) | 3.0x10<sup>-6</sup> | 1.55(1.11-2.16) | 0.01 |
|  | 4/MOD | 18 | 297 | 5.7 | 1.15(0.66-2.03) | 0.62 | 1.12(0.63-1.99) | 0.69 |
|  | 5/MARD | 76 | 466 | 14.0 | 1 | - | 1 | - |
| **Macro-albuminuria** | 1/SAID | 4 | 147 | 2.6 | 0.35(0.12-1.03) | 0.056 | 0.33(0.11-0.98) | 0.047 |
|  | 2/SIDD | 29 | 273 | 9.6 | 1.27(0.74-2.19) | 0.39 | 1.25(0.73-2.15) | 0.42 |
|  | 3/SIRD | 28 | 202 | 12.2 | 2.18(1.31-3.63) | 0.003 | 1.91(1.14-3.19) | 0.013 |
|  | 4/MOD | 27 | 276 | 8.9 | 1.31(0.72-2.38) | 0.37 | 1.34(0.74-2.45) | 0.34 |
|  | 5/MARD | 32 | 473 | 6.3 | 1 | - | 1 | - |
| **ESRD** | 1/SAID | 5 | 158 | 3.1 | 1.89(0.68-5.27) | 0.23 | 1.73(0.61-4.88) | 0.30 |
|  | 2/SIDD | 17 | 289 | 5.6 | 2.26(1.11-4.58) | 0.024 | 1.87(0.91-3.81) | 0.09 |
|  | 3/SIRD | 32 | 222 | 12.6 | 4.89(2.68-8.93) | 2.4x10<sup>-7</sup> | 3.61(1.96-6.64) | 3.9x10<sup>-5</sup> |
|  | 4/MOD | 8 | 305 | 2.6 | 1.74(0.68-4.42) | 0.25 | 1.73(0.68-4.44) | 0.25 |
|  | 5/MARD | 16 | 524 | 3.0 | 1 | - | 1 | - |

<sup>1</sup>Cox regressions adjusted for sex and age at onset.
<sup>2</sup>Cox regression adjusted for sex, age at onset and first eGFR.
CKD3A was defined as eGFR < 60 mL/min/1.73m<sup>2</sup> and (CKD 3B) as eGFR < 45 mL/min/1.73m<sup>2</sup> for more than 90 days. End-stage renal disease (ESRD) was defined as at least one eGFR below 15 mL/min/1.73m<sup>2</sup>. Diabetic kidney disease (DKD) was defined as either ESRD or macroalbuminuria defined as at least two out of three consecutive visits with albumin excretion rate (AER) ≥200 µg/min or AER ≥300 mg/24 h or albumin-creatinine ratio (ACR) ≥25/35 mg/mmol for men/women.
Median duration at first eGFR=100(IQR 54-173) days.

**Table S9.** Cox regression analysis comparing risk of diabetic retinopathy between clusters in SDR

|  |  | Events | Censored | Events (%) | HR(CI95%)<sup>1</sup> | p<sup>1</sup> | HR(CI95%)<sup>2</sup> | p<sup>2</sup> |
| --- | --- | --- | --- | --- | --- | --- | --- | --- |
| **DR** | 1/SAID | 85 | 57 | 59.9 | 0.82(0.62-1.07) | 0.82 | 1.00(0.71-1.41) | 0.99 |
|  | 2/SIDD | 169 | 96 | 63.8 | 1.25(1.01-1.55) | 0.044 | 1.49(1.17-1.89) | 0.001 |
|  | 3/SIRD | 62 | 117 | 34.6 | 0.72(0.54-0.97) | 0.029 | 0.74(0.55-1.00) | 0.048 |
|  | 4/MOD | 134 | 139 | 49.1 | 0.95(0.75-1.19) | 0.64 | 1.28(0.94-1.75) | 0.11 |
|  | 5/MARD | 165 | 252 | 39.6 | 1 | - | 1 | - |

<sup>1</sup>Cox regressions adjusted for sex.
<sup>2</sup>Cox regression adjusted for sex and age at onset.

**Table S10.** Cox regression analysis comparing risk of cardiovascular disease between clusters in SDR

|  |  | Events | Censored | Events (%) | HR(CI95%)<sup>1</sup> | p<sup>1</sup> | HR(CI95%)<sup>2</sup> | p<sup>2</sup> |
| --- | --- | --- | --- | --- | --- | --- | --- | --- |
| **CE** | 1/SAID | 20 | 138 | 12.7 | 0.43(0.27-0.69) | 4.6x10<sup>-4</sup> | 0.98(0.60-1.63) | 0.98 |
|  | 2/SIDD | 66 | 248 | 21.0 | 0.71(0.53-0.95) | 0.023 | 1.16(0.84-1.59) | 0.37 |
|  | 3/SIRD | 72 | 156 | 31.6 | 1.31(0.99-1.75) | 0.064 | 1.26(0.89-1.78) | 0.19 |
|  | 4/MOD | 52 | 262 | 16.6 | 0.56(0.41-0.78) | 0.001 | 1.43(0.83-2.45) | 0.20 |
|  | 5/MARD | 132 | 384 | 25.6 | 1 | - | 1 | - |
| **Stroke** | 1/SAID | 7 | 141 | 4.7 | 0.33(0.15-0.72) | 0.005 | 0.72(0.32-1.63) | 0.43 |
|  | 2/SIDD | 17 | 300 | 5.4 | 0.42(0.24-0.72) | 0.002 | 0.68(0.39-1.20) | 0.19 |
|  | 3/SIRD | 19 | 221 | 7.9 | 0.74(0.44-1.25) | 0.26 | 0.76(0.45-1.28) | 0.30 |
|  | 4/MOD | 14 | 285 | 4.7 | 0.32(0.18-0.57) | 1.4x10<sup>-4</sup> | 0.92(0.44-1.91) | 0.81 |
|  | 5/MARD | 56 | 484 | 10.4 | 1 | - | 1 | - |

<sup>1</sup> Cox regression adjusted for sex.
<sup>2</sup> Cox regressions adjusted for sex and age at onset.
Coronary events (CE) were defined by ICD-10 codes I21, I252, I20, I251, I253, I254, I255, I256, I257, I258, I259. Stroke was defined by ICD-10 codes I60, I61, I63 and I64.

**Table S11.**
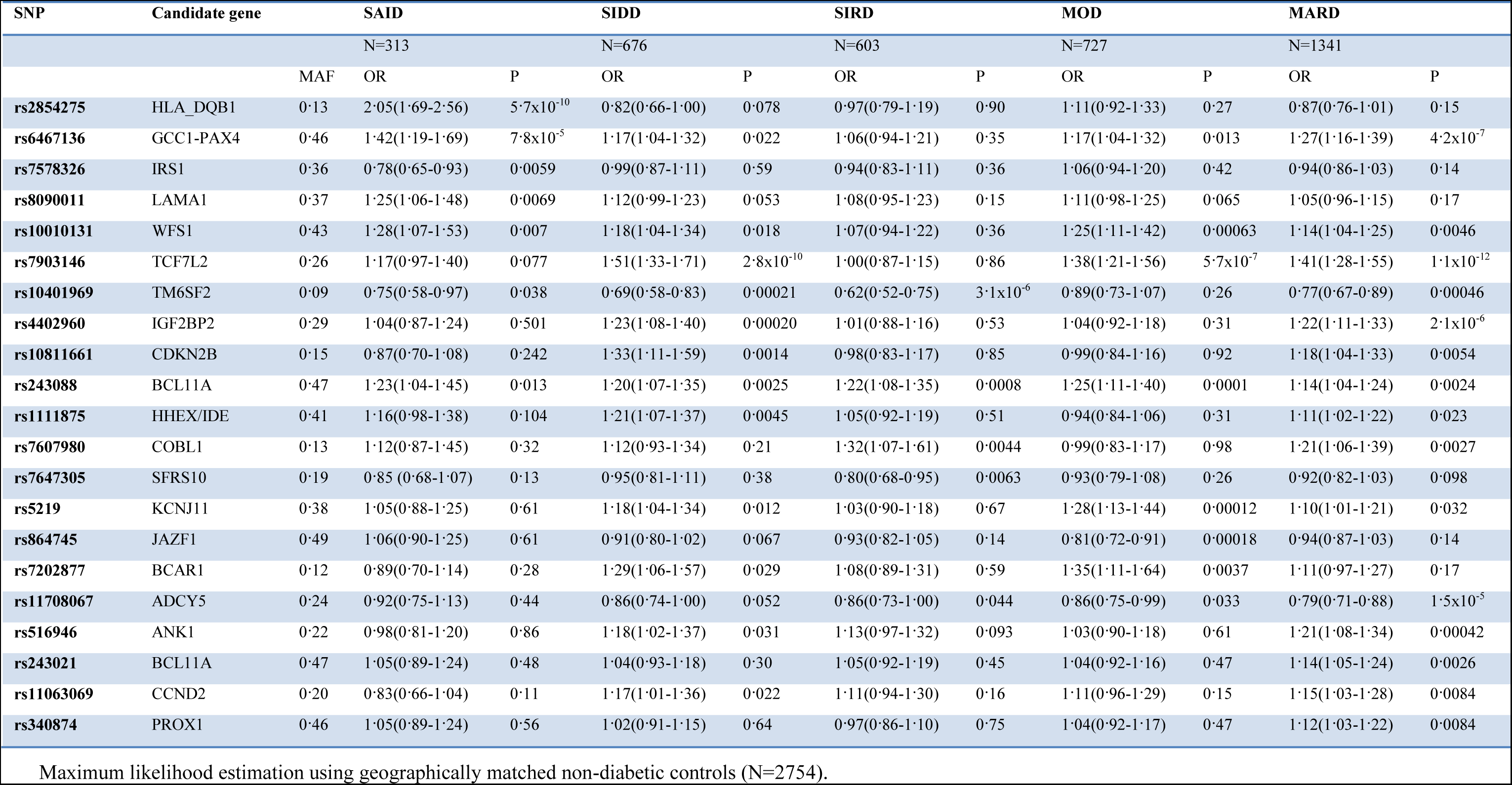
Strongest genetic associations with specific ANDIS clusters (reaching p<0.01)

## References

1. Collaboration NCDRF. Worldwide trends in diabetes since 1980: a pooled analysis of 751 population-based studies with 4.4 million participants. Lancet 2016; 387(10027): 1513–30.

2. Tuomi T, Groop LC, Zimmet PZ, Rowley MJ, Knowles W, Mackay IR. Antibodies to Glutamic Acid Decarboxylase Reveal Latent Autoimmune Diabetes Mellitus in Adults With a Non—Insulin-Dependent Onset of Disease. Diabetes 1993; 42(2): 359–62.

3. Froguel P, Zouali H, Vionnet N, et al. Familial hyperglycemia due to mutations in glucokinase. Definition of a subtype of diabetes mellitus. N Engl J Med 1993; 328(10): 697–702.

4. Yamagata K, Oda N, Kaisaki PJ, et al. Mutations in the hepatocyte nuclear factor-1alpha gene in maturity-onset diabetes of the young (MODY3). Nature 1996; 384(6608): 455–8.

5. Reddy MA, Zhang E, Natarajan R. Epigenetic mechanisms in diabetic complications and metabolic memory. Diabetologia 2015; 58(3): 443–55.

6. Brownlee M. The pathobiology of diabetic complications: a unifying mechanism. Diabetes 2005; 54(6): 1615–25.

7. Pearson ER, Flechtner I, Njolstad PR, et al. Switching from insulin to oral sulfonylureas in patients with diabetes due to Kir6.2 mutations. N Engl J Med 2006; 355(5): 467–77.

8. Lindholm E, Agardh E, Tuomi T, Groop L, Agardh CD. Classifying diabetes according to the new WHO clinical stages. Eur J Epidemiol 2001; 17(11): 983–9.

9. Manjer J, Carlsson S, Elmstahl S, et al. The Malmo Diet and Cancer Study: representativity, cancer incidence and mortality in participants and non-participants. Eur J Cancer Prev 2001; 10(6): 489–99.

10. Rahmati K, Lernmark A, Becker C, et al. A comparison of serum and EDTA plasma in the measurement of glutamic acid decarboxylase autoantibodies (GADA) and autoantibodies to islet antigen-2 (IA-2A) using the RSR radioimmunoassay (RIA) and enzyme linked immunosorbent assay (ELISA) kits. Clin Lab 2008; 54(7-8): 227–35.

11. Tuomi T, Carlsson A, Li H, et al. Clinical and genetic characteristics of type 2 diabetes with and without GAD antibodies. Diabetes 1999; 48(1): 150–7.

12. Vaziri-Sani F, Delli AJ, Elding-Larsson H, et al. A novel triple mix radiobinding assay for the three ZnT8 (ZnT8-RWQ) autoantibody variants in children with newly diagnosed diabetes. Journal of immunological methods 2011; 371(1-2): 25–37.

13. Levy JC, Matthews DR, Hermans MP. Correct homeostasis model assessment (HOMA) evaluation uses the computer program. Diabetes Care 1998; 21(12): 2191–2.

14. Levey AS, Stevens LA, Schmid CH, et al. A new equation to estimate glomerular filtration rate. Ann Intern Med 2009; 150(9): 604–12.

15. Martinell M, Dorkhan M, Stalhammar J, Storm P, Groop L, Gustavsson C. Prevalence and risk factors for diabetic retinopathy at diagnosis (DRAD) in patients recently diagnosed with type 2 diabetes (T2D) or latent autoimmune diabetes in the adult (LADA). J Diabetes Complications 2016; 30(8): 1456–61.

16. Hennig C. Cluster-wise assessment of cluster stability. Comput Stat Data An 2007; 52(1): 258–71.

17. Prasad RB, Groop L. Genetics of type 2 diabetes-pitfalls and possibilities. Genes (Basel) 2015; 6(1): 87–123.

18. Lyssenko V, Lupi R, Marchetti P, et al. Mechanisms by which common variants in the TCF7L2 gene increase risk of type 2 diabetes. J Clin Invest 2007; 117(8): 2155–63.

19. Liu YL, Reeves HL, Burt AD, et al. TM6SF2 rs58542926 influences hepatic fibrosis progression in patients with non-alcoholic fatty liver disease. Nature communications 2014; 5: 4309.

20. Li L, Cheng WY, Glicksberg BS, et al. Identification of type 2 diabetes subgroups through topological analysis of patient similarity. Science translational medicine 2015; 7(311): 311ra174.

21. Groop L, Ekstrand A, Forsblom C, et al. Insulin resistance, hypertension and microalbuminuria in patients with type 2 (non-insulin-dependent) diabetes mellitus. Diabetologia 1993; 36(7): 642–7.

22. Gnudi L, Coward RJ, Long DA. Diabetic Nephropathy: Perspective on Novel Molecular Mechanisms. Trends Endocrinol Metab 2016; 27(11): 820–30.

23. Welsh GI, Hale LJ, Eremina V, et al. Insulin signaling to the glomerular podocyte is critical for normal kidney function. Cell metabolism 2010; 12(4): 329–40.

